# Folate metabolism in tumor-associated macrophages drives immunosuppressive function to promote tumor growth

**DOI:** 10.64898/2026.09.29.755523

**Authors:** Matthew P. Zimmerman, Vasyl Zhabotynsky, Haiyang Wang, Emily K. Cox, Wan Lin Chong, Alex G. Bastian, Andrew S. Kennedy, Brian P. Fay, Amy G. Reynolds, Anna Ebacher Morrill, Jeremy A. Meier, Katherine Vietor, Khalilah E. Taylor, Pristine C. Onuoha, Siddharth Maruvada, Katie E. Hurst, Christopher J. Wong, Carlton W. Anderson, Nancy E. Thomas, Rihe Liu, Kirsten L. Bryant, Albert S. Baldwin, David W. Ollila, Jenny P-Y Ting, Blake R. Rushing, Susan J. Sumner, Sergey A. Krupenko, Willy Hugo, Stergios J. Moschos, Jessica E. Thaxton, Brian C. Miller

## Abstract

Tumor-associated macrophages (TAMs) promote tumor progression, inhibit anti-tumor immunity, and induce resistance to immune checkpoint inhibitors (ICI). Metabolism is a primary determinant of TAM polarization, suggesting therapies that selectively modulate TAM metabolic state may hold therapeutic benefit. Here we demonstrate folate uptake and metabolism sustained by the myeloid-restricted folate receptor beta (FRβ) are critical for TAM immunosuppressive function. Folate was depleted in the microenvironment of human melanomas, suggesting FRβ upregulation by TAMs may be an adaptive response to folate scarcity. FRβ⁺ TAMs exhibited an immunosuppressive phenotype and were associated with immunotherapy resistance in melanoma patients. Genetic ablation of FRβ attenuated tumor growth across multiple cancer models and repolarized macrophages toward pro-inflammatory states, promoting tumor infiltration by activated lymphocytes. Critically, excess folate reversed the phenotypes observed with FRβ loss, suggesting these effects stem from impaired folate uptake by TAMs rather than a folate-independent role for FRβ. Integrated metabolomic, transcriptomic, and functional mechanistic studies demonstrated that folate acquisition via FRβ was required to generate reduced glutathione to limit mitochondrial reactive oxygen species (ROS) production. In the absence of FRβ, mitochondrial oxidative stress promoted canonical NF-kB signaling and macrophage pro-inflammatory polarization. Thus, FRβ couples a nutrient-limited microenvironment to TAM immunosuppressive function and represents a therapeutically tractable metabolic checkpoint for TAMs.

## Introduction

Despite the success of immune checkpoint inhibitors (ICI), such as antibodies targeting PD-1/PD-L1, to treat a variety of cancers, the majority of cancer patients do not respond to these immunotherapies. Tumor-associated macrophages (TAMs) can inhibit T cell function and the efficacy of ICI and are associated with worse clinical outcomes for patients.^1–3^ However, TAM cell states are functionally and transcriptionally heterogenous, varying widely based on cancer type, developmental origin, and the local microenvironmental signals. Therapeutic strategies to deplete or repolarize immunosuppressive TAMs have had little clinical success, likely due to poor selectivity for TAMs over resident tissue macrophages (RTMs) and our limited understanding of the pathways governing TAM polarization.^4^

Cellular metabolic state is a critical determinant of macrophage polarization, particularly in the tumor microenvironment (TME), where chronic hypoxia, waste accumulation, acidic pH, and nutrient competition culminate into a metabolically-distressing locale.^5^ One carbon (1C) metabolism is a central metabolic pathway required by all somatic cells that is upregulated during both pro-inflammatory and immunosuppressive macrophage polarization.^6–9^ Folate (vitamin B9) is a required co-enzyme for 1C metabolism, and antifolate drugs that inhibit 1C metabolism are commonly used chemotherapies and immunosuppressants (e.g. methotrexate, pemetrexed, pralatrexate).^10^ In macrophages, folate deprivation or treatment with antifolates increases pro-inflammatory cytokine production, although the direct mechanisms are unknown.^11–13^

1C metabolism is critical for *de novo* nucleotide synthesis, generation of one carbon intermediates for macromolecular methylation reactions, and control of reactive oxygen species (ROS) within cells.^10^ 1C metabolism supports NADPH production, which is required to reduce the cellular antioxidant glutathione, and also regulates mitochondrial function, a significant source of ROS production.^14–17^ Consequently, folate deficiency has been demonstrated to elevate cellular ROS levels.^18^ ROS are important signaling molecules in macrophages and, depending on the intracellular localization, type, duration, and levels, can drive either pro-or anti-tumor functions.^14,15^ For example, ROS can either induce immunosuppressive macrophage polarization through activation of Keap1-Nrf2 and Jak-Stat6 signaling or, conversely, promote pro-inflammatory polarization through activation of NF-kB and HIF-1⍺ signaling.^16,17,19,20^

Myeloid cells, particularly macrophages, uniquely express a high affinity folate receptor, FRβ (gene: *Folr2*), one of 4 functional folate carriers found in mice (FRα, FRβ, RFC, PCFT; humans also have a secreted FRγ not found in mice). RFC (Reduced Folate Carrier, gene: *Slc19a1*) is expressed by all mammalian cells, whereas PCFT (Proton Coupled Folate Transporter, gene: *Slc46a1*) is involved in folate uptake in low pH environments (e.g. gut lumen, choroid plexus).^21^ Unlike these low-affinity folate transporters, FRβ has about 1000x higher affinity for folates, particularly oxidized folates such as folic acid.^21^ However, because it is GPI anchored, folates bound to FRβ must enter the cell through endocytosis, instead of direct membrane transport, resulting in lower capacity uptake than the folate transporters.^22^ Unlike mice with knockouts of the other folate transporters, *Folr2*^-/-^ mice are viable and do not have neural tube defects.^21,23^ FRβ is upregulated on activated macrophages and TAMs in the TME, where it has been associated with both immunosuppressive and pro-inflammatory phenotypes.^24–28^ Recently, it has been suggested that FRβ may have functions independent of folate uptake – regulating macrophage migration, serving as a direct T cell checkpoint, promoting inflammasome activation, and sustaining TGF-β signaling.^29–32^ Given its upregulation on TAMs, multiple groups have developed FRβ-targeted therapeutics to deplete FRβ^+^ TAMs or deliver specific drugs to these cells,^33–38^ however we still do not understand how FRβ regulates TAM biology nor the role of FRβ^+^ TAMs in mediating ICI resistance.

Here we demonstrate that FRβ marks immunosuppressive TAMs and predicts poor outcomes in melanoma patients, raising the question of whether this receptor is a passive biomarker or an active requirement for TAM-mediated immunosuppression. To interrogate this question, we used CRISPR-Cas9 to generate novel *Folr2*^-/-^ mice on the C57BL/6J background. Tumors implanted into *Folr2*^-/-^ mice grew more slowly, with increased anti-tumor TAM polarization and activation of innate and adaptive lymphocytes. Consequently, loss of FRβ synergized with anti-PD-1 treatment to better control tumor growth. To study the functions of FRβ, we cultured *Folr2*^-/-^ bone marrow-derived macrophages (BMDMs) in media with a physiologic concentration of folate and determined these cells replicated the transcriptional and functional phenotypes of *Folr2*^-/-^ TAMs observed *in vivo*. These phenotypes were due to deficient folate uptake by *Folr2*^-/-^ cells: feeding *Folr2*^-/-^ mice a high-folate diet or growing *Folr2*^-/-^ BMDMs in supraphysiologic-folate media reversed tumor growth phenotypes and pro-inflammatory macrophage polarization. Mechanistically, FRβ-mediated folate uptake was required for regulation of mitochondrial function and ROS production. In the absence of FRβ, oxidative stress promoted mitochondrial dysfunction, activated NF-κB signaling, and induced macrophage pro-inflammatory polarization. Together, our data demonstrate that TAMs require FRβ for folate uptake in the low-folate TME to prevent mitochondria-induced pro-inflammatory polarization.

## Results

### FRβ is expressed by immunosuppressive TAMs and is associated with poor outcomes for patients with melanoma

FRβ expression has been associated with both immunosuppressive and pro-inflammatory macrophages, depending on the specific tumor type or inflammatory context.^24–28^ To evaluate the role of FRβ^+^ TAMs in melanoma, we integrated and analyzed two published single-cell RNA-seq (scRNA-seq) datasets from melanoma patients.^39,40^ We identified a robust population of *FOLR2*^+^ TAMs that co-expressed the pro-tumor TAM markers *MRC1* (CD206), *CD209*, and *CD163* (Fig. 1A). Next, using fresh biopsies of 6 tumors obtained from 4 patients with advanced melanoma, we demonstrated approximately 60% of TAMs expressed FRβ in advanced tumors (Fig. 1B, Supp. Fig. 1A). FRβ^+^ TAMs had higher expression of the immunosuppressive markers CD163, CD206, and PD-L1 versus FRβ^-^ TAMs, supporting our previous scRNA-seq analysis (Fig. 1C, Supp. Fig. 1B). To determine whether FRβ⁺ TAM abundance is associated with overall survival in melanoma, we stratified patient survival in the melanoma (SKCM) TCGA dataset (n = 432 patients, most tumors collected before ICI approval) based on the relative expression of *FOLR2* and *CD8A*. Patients with a high ratio of *FOLR2/CD8A* – suggesting a relative abundance of FRβ^+^ TAMs over CD8^+^ T cells – had significantly worse overall survival (Fig. 1D). Survival analysis using a Cox model adjusting for *CD8A*, age, sex, and weight also showed that higher expression of *FOLR2* was associated with worse survival (Supp. Fig. 1C). Together, these results suggest that FRβ^+^ TAMs possess an immunosuppressive phenotype and are associated with worse outcomes in melanoma patients.

**Figure 1.**
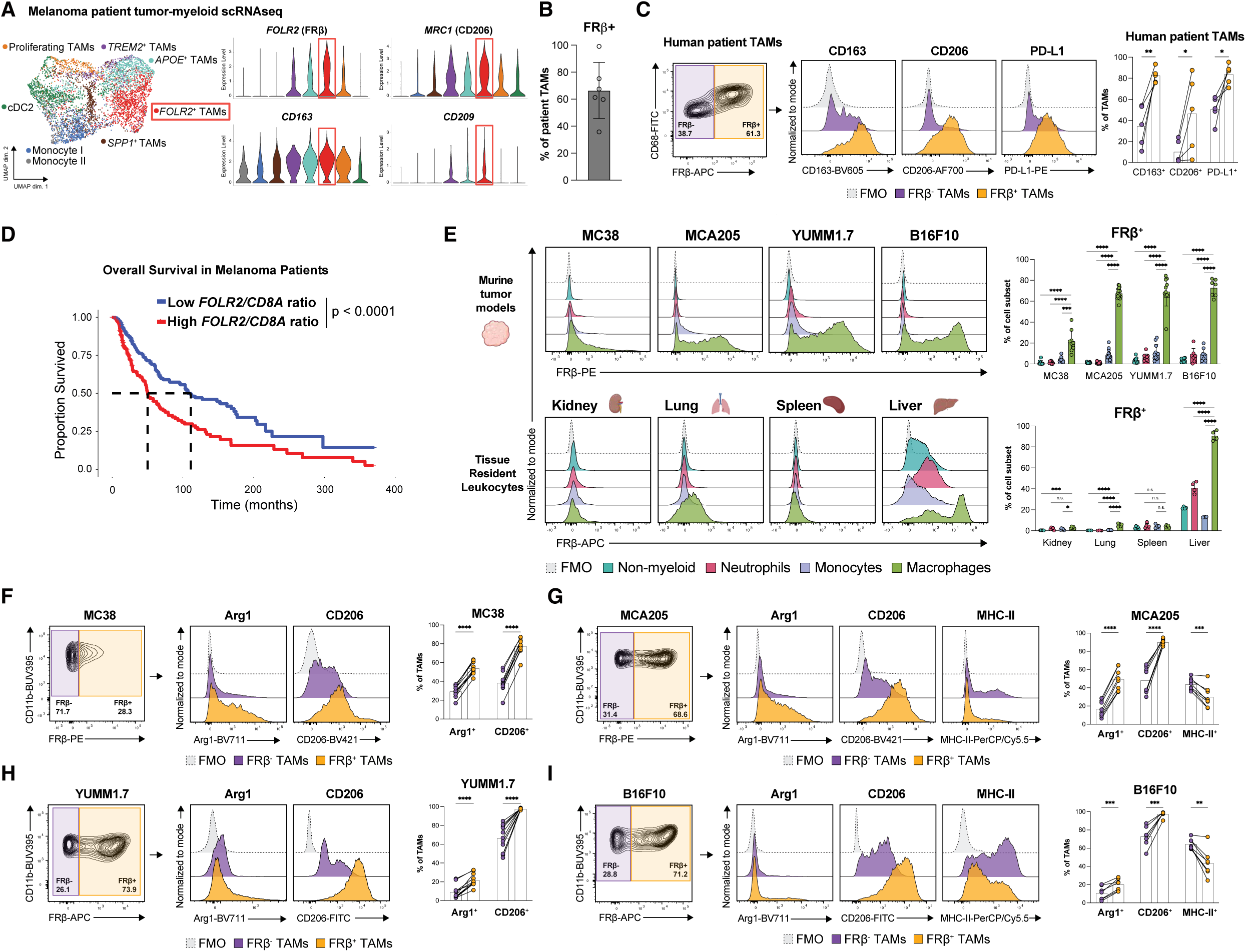
FRβ is expressed by TAMs with an immunosuppressive phenotype and is associated with poor melanoma patient outcomes. **(A)** Single-cell data combined from Sade-Feldman *et al.* and Pozniak *et al.* and subclustered on myeloid cells. *FOLR2*^+^ TAM cluster shown in red; co-expression of *FOLR2* and markers of immunosuppressive macrophages (*MRC1*, *CD163*, *CD209*) shown. **(B)** Percentage of human TAMs expressing FRβ from 6 melanoma tumors isolated from 4 patients undergoing metastatic melanoma resection for Lifileucel treatment. **(C)** Flow cytometry analysis of FRβ^+^ and FRβ^-^ human TAMs from 5 patient tumors showing expression of CD163, CD206, and PD-L1 in each population. Representative flow plots (left) and summary bar plots (right) from 5 patient tumors (excluding n=1 tumor with 100% FRβ^+^ TAMs). **(D)** Kaplan-Meier estimate of survival in the melanoma (SKCM) cohort of TCGA, stratified as *FOLR2/CD8A* ratio high or low using 60/40 percentile as cutoff value. At all cutoffs between 0.2 and 0.8 the p-value from log-rank test was significant. **(E)** Flow cytometry analysis of FRβ expression by mouse tumor-infiltrating and tissue resident leukocytes across 4 subcutaneous tumor models and 4 organs. Representative histograms (left) and summary bar plots (right) shown. All tumor data pooled from two independent experiments. Peripheral organ analysis from one of two representative independent experiments. **(F-I)** Flow cytometry analysis of FRβ^+^ and FRβ^-^ TAMs from MC38 **(F)**, MCA205 **(G)**, YUMM1.7 **(H)** and B16F10 **(I)** tumor models showing expression of Arg1, CD206, and MHC-II. Representative histograms (left) and summary bar plots from 2 pooled experiments shown (1 experiment for MCA205). * p < 0.05; ** p <0.01; *** p < 0.001; **** p < 0.0001. Created in BioRender. Zimmerman, M. (2027) https://BioRender.com/wk9olvt

Next, we evaluated the expression of FRβ by leukocytes in both the TME and peripheral organs (kidney, lung, spleen, liver) in mice. We observed between 20% - 80% of TAMs expressed FRβ, depending on the specific tumor model (Fig. 1E, Supp. Fig. 1D). Importantly, this expression was primarily limited to TAMs and not robustly observed on any other cell type in the tumor or peripheral organs, with the exception of the liver, where FRβ has been reported as a canonical marker of Kupffer cells (Fig. 1E, Supp. Fig. 1E).^32,41^ In the Human Protein Atlas, *Folr2* transcript and FRβ protein were only reliably detected in macrophages (Supp. Fig. 1F).^42,43^ Finally, similar to our melanoma patient analysis, we discovered FRβ^+^ TAMs had higher expression of Arginase-1 (Arg1) and CD206 across all tumor models as well as lower expression of MHC-II in some models compared with FRβ^-^ TAMs, consistent with a more immunosuppressive phenotype (Fig. 1F-I, Supp. Fig. 1G).

### FRβ is required for immunosuppressive TAM polarization *in vivo*

To evaluate FRβ’s mechanistic role in TAM polarization and anti-tumor immunity, we generated *Folr2*^-/-^ mice on the C57BL/6J background using CRISPR-Cas9 to remove all coding DNA sequences (Fig. 2A). We observed no discernible birth defects or deviations from expected pup numbers (data not shown). BMDMs and TAMs isolated from *Folr2*^-/-^ mice completely lacked *Folr2* transcript and FRβ protein expression (Fig. 2B, 2C).

**Figure 2.**
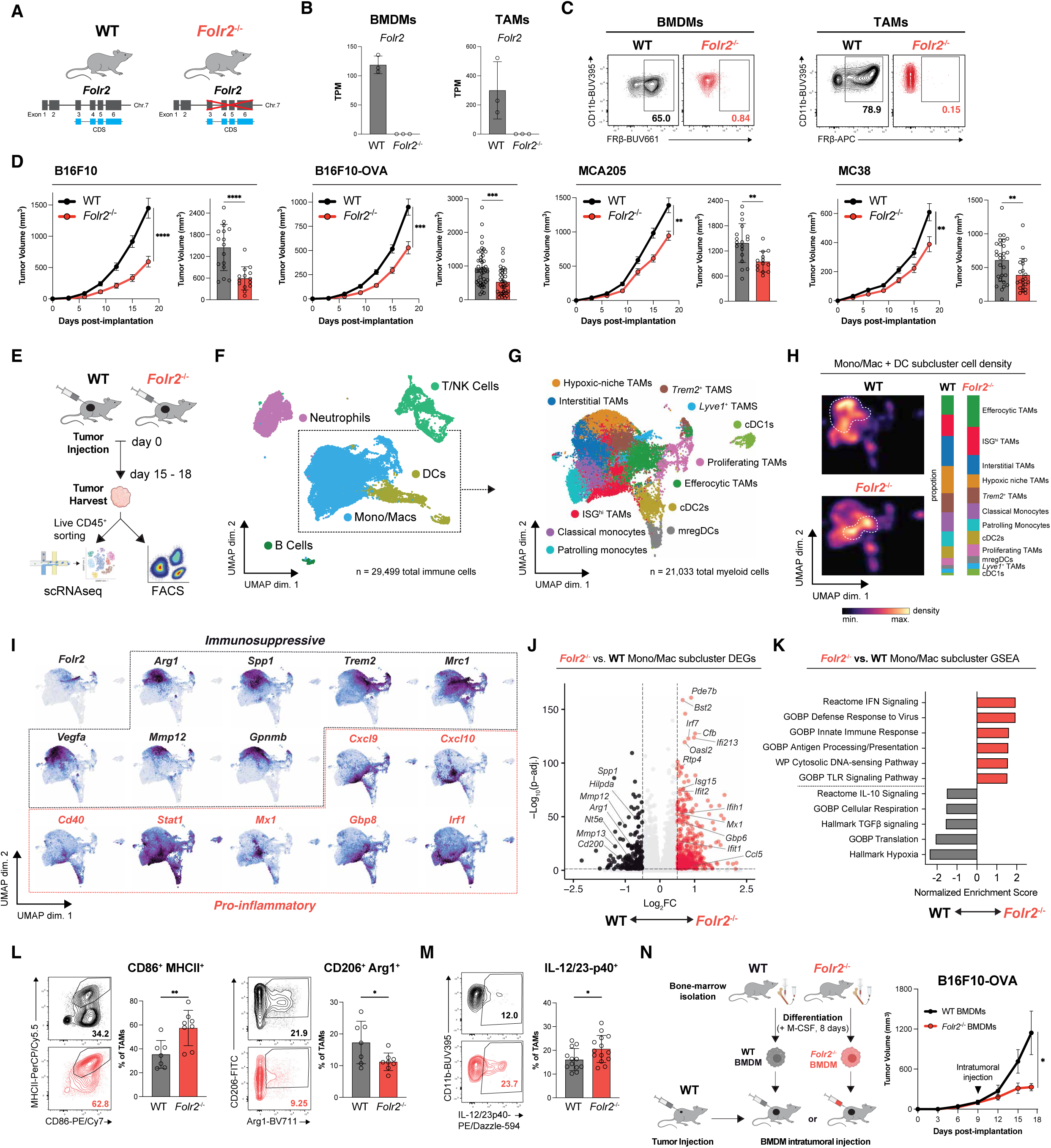
FRβ is critical for TAM immunosuppressive polarization and function to promote tumor growth. **(A)** Schema of *Folr2*^-/-^ mouse generation. **(B)** Bulk RNA-seq analysis of *Folr2* expression in bone marrow-derived macrophages (BMDMs) or TAMs sorted from B16F10-OVA tumors isolated from wild-type or *Folr2^-/-^*mice. **(C)** Flow cytometry analysis of FRβ expression in wild-type and *Folr2*^-/-^ BMDMs and TAMs isolated from B16F10 tumors. **(D)** Tumor growth data for the indicated tumor cell lines implanted into wild-type or *Folr2*^-/-^ mice, pooled from 3 experiments (B16F10), 5 experiments (B16F10-OVA), 2 experiments (MCA205), and 4 experiments (MC38). **(E)** Schema of wild-type and *Folr2*^-/-^ mouse tumor immunophenotyping. **(F, G)** UMAPs of tumor-infiltrating total leukocytes **(F)** and Mono/Mac and dendritic cell (DC) clusters **(G)** from B16F10 tumors implanted into n =5 wild-type or n=5 *Folr2*^-/-^ mice. **(H)** Mono/Mac and DC density via galaxy plot in UMAP space (left) and quantification of the proportion of cells in each cluster between wild-type and *Folr2*^-/-^mice (right). Pooled from 5 tumors implanted in each genotype of mice. **(I)** Expression of key genes in individual myeloid cells from B16F10 tumors. **(J)** Volcano plot of differential gene expression between wild-type and *Folr2*^-/-^ Mono/Mac cluster cells. *Folr2* omitted from the plot as its log₂ fold change (-5.0) falls outside the displayed x-axis range. **(K)** Gene set enrichment analysis (GSEA) of indicated signatures in the ranked list of genes differentially expressed between all Mono/Mac cluster cells isolated from wild-type and *Folr2*^-/-^ mice. **(L)** Flow cytometry analysis of wild-type and *Folr2*^-/-^ TAMs isolated from B16F10 tumors stained for CD86, MHC-II, CD206, and Arg1. Representative flow plots and pooled bar plots from 2 independent experiments shown. **(M)** Frequency of IL-12/23p40^+^ TAMs isolated from wild-type or *Folr2*^-/-^ mice and cultured for 18h *ex vivo*. Representative contour flow plots (left) and quantification (right) pooled between three independent experiments. **(N)** 1×10^6^ wild-type or *Folr2*^-/-^ BMDMs cultured in physiologic-folate media without polarizing cytokines were injected intratumorally on day 9 into established B16F10-OVA tumors implanted into wild-type mice. Tumor growth measured. One of 3 independent experiments shown. * p < 0.05; ** p < 0.01; *** p < 0.001; **** p < 0.0001; ns not significant. Created in BioRender. Zimmerman, M. (2027) https://BioRender.com/kn294bd, https://BioRender.com/t4gkhma

Tumor growth was significantly decreased in *Folr2*^-/-^ mice across multiple syngeneic tumor models, with a 60% reduction in parental B16F10 tumors and 45% reduction in immunogenic B16F10-OVA tumors (Fig. 2D). We next isolated CD45^+^ tumor-infiltrating leukocytes from B16F10 tumors and performed scRNA-seq and flow cytometry analysis to interrogate how loss of *Folr2* globally altered the TME (Fig. 2E, Supp. Fig. 2A-C). First, we confirmed that the *Folr2* transcript is only detected in monocyte/macrophage clusters (Mono/Macs) (Supp. Fig. 2D, 2E). The TME of *Folr2*^-/-^ mice was depleted of interstitial (*Msr1*, *Vcan*, *Spp1*) and hypoxic-niche (*Vegfa*, *Hif1a*, *Egln3*) TAMs and enriched in ISG^hi^ (*Mx1*, *Isg15*, *Cxcl10*) and efferocytic (*Axl*, *Mertk*) TAMs, suggesting a dramatic repolarization of the myeloid compartment (Fig. 2F-H, Supp. Fig. 2F, 2G). Indeed, differential gene expression analysis of Mono/Mac clusters from *Folr2*^-/-^ and control mice revealed enrichment of pro-inflammatory gene expression in *Folr2*^-/-^ cells, marked by interferon (IFN) signaling, antigen presentation machinery, and innate immune response pathways (Fig. 2I-K). Conversely, wild-type Mono/Macs were characterized by immunosuppressive gene expression, dominated by TGF-β signaling, IL-10 signaling, and hypoxia response pathways (Fig. 2I-K). These findings were orthogonally confirmed by an enrichment of pro-inflammatory gene expression pathways in bulk RNAseq analysis of wild-type and *Folr2*^-/-^ TAMs flow-sorted from B16F10-OVA tumors (Supp. Fig. 2H, Supp. Table 1). In addition to our Mono/Mac analysis, we also observed an enrichment of interferon-response and other pro-inflammatory gene signatures in neutrophils in *Folr2*^-/-^ tumors (Supp. Fig. 2I-M). Given neutrophils completely lacked *Folr2* expression, these findings were consistent with these cells responding to a more inflamed TME as opposed to a cell-intrinsic effect of *Folr2* deletion (Supp. Fig. 2D, 2E, 2N). Pseudotime analysis of neutrophils demonstrated that these cells branched early and progressed down a distinct differentiation pathway in *Folr2*^-/-^ mice, likely driven by interferon signaling (Supp. Fig. 2O). Confirming our transcriptional analysis, flow cytometry of TAMs from B16F10 tumors revealed an increase in CD86 and MHC-II expression and a decrease in CD206 and Arg1 expression, consistent with a pro-inflammatory TAM phenotype (Fig. 2L). To functionally evaluate these TAMs, we sorted myeloid cells from B16F10 tumors for *ex vivo* cytokine production and found that *Folr2*^-/-^ TAMs made more pro-inflammatory IL-12/23p40 (Fig. 2M). Finally, to directly test if *Folr2*^-/-^ macrophages control tumor growth, we cultured BMDMs from wild-type or *Folr2*^-/-^ mice and injected these cells into established B16F10-OVA tumors on day 9 post tumor implantation. *Folr2*^-/-^ BMDMs significantly reduced tumor growth compared with control BMDMs (Fig. 2N). Together, these results demonstrate that FRβ promotes TAM immunosuppressive polarization in the TME.

### FRβ^+^ TAMs inhibit immune effector cells

To evaluate the function of FRβ^+^ TAMs in human melanoma, we first performed spatial transcriptomics of liver, lung, lymph node, and brain melanoma metastases isolated from 8 patients who donated their bodies to the UNC warm autopsy program.^44,45^ Because our Visium data does not have single-cell resolution, instead capturing up to 10 cells per spatial cluster, we generated cell-type signatures from scRNA-seq data and used these to deconvolute cell populations within Visium clusters. We identified many clusters enriched in *FOLR2^+^* TAMs (Fig. 3A, red clusters). *FOLR2^+^* TAM clusters frequently colocalized with exhausted CD8^+^ T cells but had moderate to poor spatial overlap with other T cell activation states (Fig. 3B). This analysis supports the hypothesis that FRβ^+^ TAMs inhibit T cell activation and promote dysfunction.

**Figure 3.**
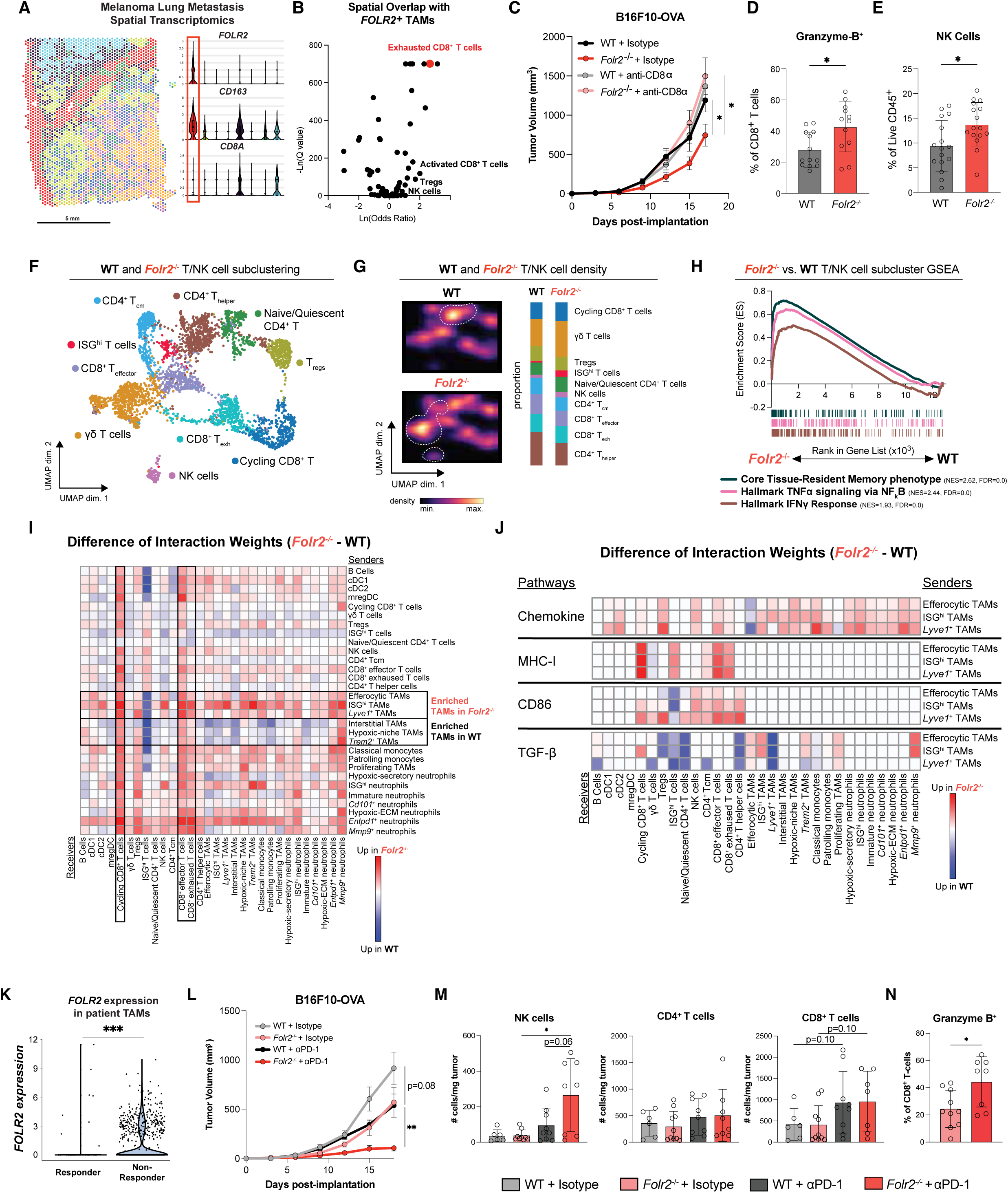
*Folr2*^-/-^ TAMs promote tumor-infiltrating lymphocyte activation, anti-tumor immunity, and immune checkpoint inhibitor efficacy. **(A)** Visium spatial transcriptomic analysis of a melanoma lung metastasis sample, demonstrating a cluster containing *FOLR2^+^* TAMs (red). **(B)** Volcano plot of spatial overlap (Fisher’s exact test) of *FOLR2^+^* TAMs with different immune cells from 22 melanoma resections across 4 metastatic sites. **(C)** B16F10-OVA tumor volumes from wild-type or *Folr2*^-/-^ mice treated with anti-CD8α depleting or isotype control antibodies. Results pooled from three independent experiments. **(D)** Quantification of Granzyme B-expressing CD8^+^ T cells from B16F10-OVA tumors in wild-type and *Folr2*^-/-^ mice. **(E)** Quantification of NK cell frequency in B16F10 tumors in wild-type and *Folr2*^-/-^ mice. **(F)** scRNA-seq profiles of tumor-infiltrating lymphocytes cells from B16F10 tumors implanted into wild-type or *Folr2*^-/-^ mice. UMAP of major T/NK cell clusters shown. **(G)** T/NK cell density via galaxy plot in UMAP space (left) and quantification of the proportion of cells in each cluster between wild-type and *Folr2*^-/-^ mice (right). Pooled from 5 tumors implanted in each genotype of mice. **(H)** GSEA of indicated signatures in T/NK cells from B16F10 tumors. Core Tissue Resident Memory phenotype signature adapted from Milner *et al*. **(I)** Heatmap showing difference of weighted interactions (*Folr2*^-/-^ minus wild-type) between single-cell transcriptional clusters from CellChat analysis. Positive scores (red) indicate higher interaction between cells from *Folr2*^-/-^ mice, negative scores (blue) indicate higher interaction between cells from wild-type mice. **(J)** Heatmap showing difference of weighted interactions (*Folr2*^-/-^ minus wild-type) for the specific pathways and sender populations shown. **(K)** ScRNA-seq of *FOLR2* expression in TAMs from ICI-responder and non-responder melanoma patients from Sade-Feldman *et al*. **(L)** Growth curves of B16F10-OVA tumors implanted into wild-type or *Folr2*^-/-^ mice treated with either anti-PD-1 (αPD-1) or isotype control antibodies every 3 days starting once tumors reached an average of 100mm^3^. Pooled results from two independent experiments. **(M)** Quantification of NK, CD4^+^, and CD8^+^ T cell infiltration into tumors from the four treatment conditions shown in **(L)**. **(N)** Quantification of Granzyme B-expressing CD8^+^ T cells in *Folr2*^-/-^ mice receiving αPD-1 or isotype control antibodies. * p < 0.05; ** p < 0.01; *** p < 0.001; **** p < 0.0001; ns not significant.

To determine if CD8^+^ T cells are required for control of B16F10-OVA tumors in *Folr2*^-/-^ mice, we depleted >98% of CD8^+^ T cells from the TME using anti-CD8α antibodies (Supp Fig. 3A, 3B). In the absence of CD8^+^ T cells, there was no longer a difference in tumor size between *Folr2*^-/-^ and control mice, suggesting the anti-tumor effects of *Folr2* loss are primarily dependent on improved CD8^+^ T cell-mediated tumor control (Fig. 3C). Indeed, we confirmed an increase in tumor-infiltrating Granzyme B^+^ CD8^+^ T cells in *Folr2*^-/-^ mice (Fig. 3D). However, we also observed tumor control in cancer models without strong xenoantigens, such as B16F10. We did not detect major changes in the numbers or activation states of CD8^+^ T cells or CD4^+^ T cells in B16F10 tumors by flow cytometry (data not shown). However, NK cell numbers were increased in B16F10 tumors in *Folr2*^-/-^ mice compared with control mice, suggesting activation of other innate lymphocytes to control B16F10 tumor growth (Fig. 3E). Returning to our scRNA-seq data from B16F10 tumors, we observed an increase in innate-like lymphocytes such as yδ T cells and NK cells (Fig. 3F, 3G, Supp. Fig. 3C). Gene set enrichment analysis (GSEA) of T/NK cells revealed an enrichment of tissue-resident memory phenotype and inflammatory response pathway gene signatures in lymphocytes from *Folr2*^-/-^ tumors (Fig. 3H).^46^ These data support the hypothesis that FRβ expression on TAMs is required for inhibition of multiple immune effectors, depending on the specific tumor context.

To further interrogate the potential crosstalk between myeloid cells and TILs, we performed CellChat, an algorithm that predicts potential interactions between cell populations using known ligand-receptor pairs.^47^ We observed a general increase in communication between the efferocytic, ISG^hi^, and *Lyve1^+^*(*Lyve1, Selenop, Cd163*) TAMs with myeloid cells, cycling lymphocytes, and effector/exhausted CD8^+^ T cells in *Folr2*^-/-^ mice (Fig. 3I, Supp. Fig. 3D). The most highly upregulated receptor-ligand pairs identified in these TAM populations were MHC-I, CD86, and chemokine pathways while the most consistently downregulated was TGF-β signaling, in agreement with with the transcriptional decrease in this pathway we previously observed (Fig. 3J, 2K). Conversely, signaling from TAMs with an immunosuppressive phenotype (interstitial, hypoxic-niche, and *Trem2*^+^) was globally decreased in the absence of *Folr2* (Fig. 3I, Supp. Fig. 3D). These findings suggest an upregulation of antigen presentation and immune cell activation and recruitment as well as downregulation of immunosuppressive signaling by *Folr2*^-/-^ TAMs.

Given the increase in predicted immunostimulatory interactions between myeloid and TIL populations in *Folr2*^-/-^ tumors, we hypothesized that ablation of *Folr2* could increase the therapeutic efficacy of anti-PD-1 treatment. Using a published single-cell dataset of 32 melanoma patients who received ICI,^40^ we identified an increase in *FOLR2*^+^ TAMs in non-responding patients (Fig. 3K). Next, we asked if anti-PD-1 immunotherapy is more effective in *Folr2*^-/-^ mice. We observed a dramatic decrease in B16F10-OVA tumor growth in *Folr2*^-/-^ mice treated with anti-PD-1 compared with anti-PD-1 treated wild-type mice (Fig. 3L). Flow cytometry revealed an increase in immune effector cells (NK cells and CD8^+^ T cells) and Granzyme B^+^ CD8^+^ T cells in anti-PD-1 treated compared with isotype-treated *Folr2*^-/-^ mice (Fig. 3M, 3N). Together, these results suggest that repolarizing TAMs into pro-inflammatory cells by loss of *Folr2* promotes TIL activation and improves the efficacy of anti-PD-1 therapy.

### Decreased folate uptake is responsible for the pro-inflammatory phenotype of *Folr2*^-/-^ macrophages

Given that FRβ has been postulated to have folate-independent functions in addition to its role as a folate carrier,^29–32^ we wanted to test if the phenotype of *Folr2*^-/-^ TAMs is due to deficient folate uptake. First, we assessed folate species in untargeted metabolomics of melanoma metastases and adjacent healthy tissue isolated from 14 warm autopsy patients.^44,45^ We observed a decrease in intratumoral folate species relative to adjacent normal tissue, suggesting melanoma tumors are depleted of folate (Fig. 4A). To model the low folate TME *in vitro*, we directly quantified folate levels in tumor interstitial fluid (TIF) and found that TIF folate concentrations ranged from 5 - 10ng/mL in both mouse and human tumors, 100 - 200-fold less than the 1000ng/mL found in standard tissue culture media (Fig. 4B). Given prior studies suggesting IL-4-treated BMDMs most faithfully recapitulate the differentiation state of TAMs *in vivo*,^48^ BMDMs were cultured in physiologic (10ng/mL) or supraphysiologic (1000ng/mL) concentrations of folic acid and polarized with IL-4. In physiologic-folate conditions, we detected a significant decrease in the intracellular folate concentration in *Folr2*^-/-^ BMDMs compared with wild-type BMDMs (Fig. 4C). However, differences in intracellular folate were rescued by supraphysiologic-folate levels (Fig. 4C). While both TAMs and BMDMs express the low-affinity folate transporters *Slc19a1* and *Slca46a1*, *Folr2* was expressed at >10x higher levels than either of these folate transporters (Fig. 4D, Supp. Fig. 4A, 4B). Therefore, we hypothesized that at sufficiently high folate concentrations, low expression of the folate transporters can compensate for the loss of FRβ to maintain intracellular folate levels, but FRβ is critically required in low-folate environments.

**Figure 4.**
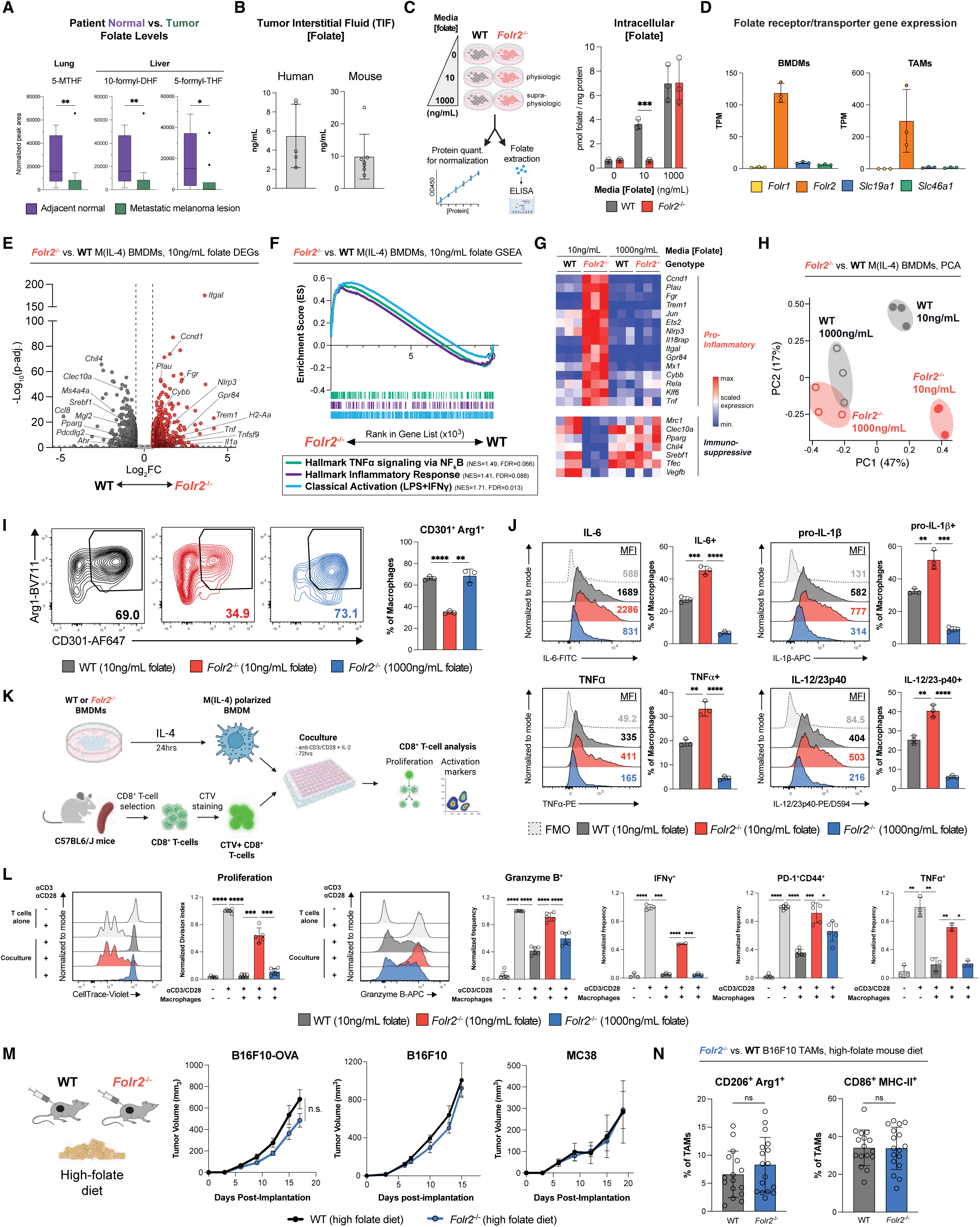
Folate uptake by FRβ is required for immunosuppressive macrophage polarization and function. **(A)** Folate metabolites assessed from untargeted metabolomics performed on human metastatic melanoma samples and adjacent normal lung and liver. **(B)** Measurement of tumor interstitial fluid (TIF) from human and mouse tumors by folate ELISA. **(C)** Intracellular folate measurements by ELISA of BMDMs grown for 6 days in folate-free media + dialyzed FBS supplemented with 0, 10, or 1000ng/mL folic acid normalized to protein quantification. **(D)** Folate receptor and transporter expression in bulk RNA-seq data from TAMs flow-sorted from B16F10-OVA tumors or M(IL-4) BMDMs grown in physiologic-folate media. Transcripts per million (TPM) shown. **(E)** Volcano plot of differentially expressed genes between wild-type and *Folr2*^-/-^ M(IL-4) BMDMs cultured in physiologic folate; *Folr2* omitted from the plot as its log₂ fold change (-11.8) falls outside the displayed x-axis range. **(F)** GSEA of indicated signatures in the ranked list of genes differentially expressed between wild-type and *Folr2*^-/-^ M(IL-4) BMDMs grown in physiologic-folate media. **(G)** Heatmap of representative pro-inflammatory and immunosuppressive gene expression in *Folr2*^-/-^ M(IL-4) BMDMs cultured in either physiologic- or supraphysiologic-folate media. **(H)** PCA of bulk RNA-seq data from wild-type and *Folr2*^-/-^ M(IL-4) BMDMs grown in physiologic- or supraphysiologic-folate media. **(I)** Representative flow plots (left) and quantification (right) of CD301 and Arg1 co-expression in wild-type and *Folr2*^-/-^ M(IL-4) BMDMs. 1 of 2 independent experiments shown. **(J)** Representative histograms and quantification of intracellular cytokine staining in wild-type and *Folr2^-/-^* M(IL-4) BMDMs. 1 of 3 independent experiments shown. **(K)** Schema of *in vitro* CD8^+^ T cell inhibition assay with BMDMs. **(L)** Representative flow plots and quantification of CellTrace Violet dilution and expression of Granzyme B, PD-1^+^CD44^+^, IFNу, and TNFα by flow cytometry of CD8^+^ T cells co-cultured with BMDMs at a 1:4 or 1:8 (BMDM:T cell) ratio. Pooled data from two of three or representative data from one of three independent experiments shown. Data normalized to stimulated T cell condition without macrophages. **(M)** Tumor growth curves of B16F10-OVA, B16F10, and MC38 in wild-type and *Folr2*^-/-^ fed high folate diet for 4 weeks prior to tumor implantation. Data shown from 2-3 pooled or 1 of 2 representative experiments. **(N)** Flow cytometry analysis of wild-type and *Folr2*^-/-^ TAMs isolated from B16F10 tumors implanted in mice fed a high-folate diet stained for CD86, MHC-II, CD206, and Arg1. Data pooled from 2 independent experiments shown. * p < 0.05; ** p < 0.01; *** p < 0.001; **** p < 0.0001; ns not significant. Created in BioRender. Zimmerman, M. (2027) https://BioRender.com/gpx6uoy, https://BioRender.com/a4m6ekl, https://BioRender.com/2bolyem

Consistent with our *Folr2*^-/-^ TAM scRNA-seq and bulk RNA-seq analyses, we observed an enrichment of pro-inflammatory gene expression in *Folr2*^-/-^ BMDMs (Fig. 4E, 4F, Supp. Table 2,3). There was excellent overlap between signatures enriched in *Folr2*^-/-^ TAMs and IL-4-polarized BMDMs, suggesting shared biology mediated by loss of FRβ both *in vitro* and *in vivo* (Supp. Fig. 4C). In addition to increased pro-inflammatory gene expression, we also found decreased immunosuppressive gene expression in *Folr2*^-/-^ BMDMs (Fig. 4E, 4G). Interestingly, the changes in expression of pro-inflammatory and immunosuppressive genes could be reversed by culturing *Folr2*^-/-^ BMDMs in supraphysiologic-folate media (Fig. 4G). Principle component analysis (PCA) revealed that the dramatic transcriptional differences between wild-type and *Folr2*^-/-^ macrophages were eliminated when the cells were cultured in supraphysiologic-folate media (Fig. 4H). These data support the hypothesis that FRβ-mediated folate uptake is critical to promote immunosuppressive macrophage polarization.

We next utilized our *in vitro* BMDM system to comprehensively evaluate how *Folr2* deletion influenced macrophage function. We noted a decrease in *Folr2*^-/-^ BMDM numbers in physiologic-folate media due to an increase in cell death (Supp. Fig. 4D-F), therefore we either normalized results for cell number or performed single-cell readouts for all subsequent functional studies. Co-expression of CD301 and Arg1, canonical markers of immunosuppressive polarization, was decreased in *Folr2*^-/-^ BMDMs, but was rescued by supraphysiologic-folate media (Fig. 4I). Similarly, production of pro-inflammatory cytokines IL-12/23p40, TNFα, pro-IL-1β, and IL-6 was increased in *Folr2*^-/-^ BMDMs and reversed by supraphysiologic-folate media (Fig. 4J). We next tested if *Folr2*^-/-^ BMDMs could inhibit T cell activation by co-culturing *Folr2*^-/-^ and wild-type BMDMs with CTV-labeled CD8^+^ T cells stimulated with anti-CD3/CD28 (Fig. 4K). While wild-type BMDMs robustly inhibited T cell proliferation, cytokine production, and activation, *Folr2*^-/-^ BMDMs cultured in physiologic-folate media did not (Fig. 4L). *Folr2*^-/-^ BMDMs cultured in supraphysiologic-folate media reversed this phenotype (Fig. 4L, Supp. Fig. 4G). Finally, we fed mice a high-folate diet (8.6mg/kg, compared with 0.2mg/kg low-folate diets) that increased their serum folate levels 8- to 10-fold (Supp. Fig. 4H). Tumor growth in wild-type mice was unchanged when comparing mice on low- and high-folate diets, suggesting that the different diets did not directly impact tumor cell growth (Supp. Fig. 4I). However, *Folr2*^-/-^ mice fed a high-folate diet no longer controlled tumor growth better than control mice, arguing sufficient levels of dietary folate could rescue the loss of FRβ on TAMs *in vivo* (Fig. 4M). The pro-inflammatory phenotype of *Folr2*^-/-^ TAMs was also lost in mice fed a high-folate diet (Fig. 2L, 4N). Taken together, our *in vitro* and *in vivo* rescue data strongly support the conclusion that the primary function of FRβ on TAMs is to maintain folate uptake in low-folate environments to support immunosuppressive polarization.

### Folate uptake is required to prevent mitochondrial oxidative stress in macrophages

To interrogate the mechanisms by which impaired folate uptake due to loss of FRβ induces pro-inflammatory polarization, we performed untargeted metabolomics on *Folr2*^-/-^ and wild-type BMDMs cultured in physiologic- or supraphysiologic-folate media and integrated this analysis with our transcriptional data (Supp. Fig. 5A, Supp. Table 4). Analogous to our bulk RNA-seq PCA results, sparse partial least squares-discriminant analysis (sPLS-DA) revealed prominent metabolomic differences between *Folr2*^-/-^ and wild-type BMDMs cultured in physiologic-folate media that were lost in supraphysiologic conditions (Fig. 5A, Supp. Fig. 5B). Metabolomic pathway enrichment analysis identified expected changes in one carbon (1C) metabolism and related pyrimidine, glutathione, and glycine/serine metabolic pathways (Fig. 5B). Reassuringly, folate species and associated catabolites (5-methyltetrahydrofolate, folic acid, and pterine) were the most highly depleted metabolites in *Folr2*^-/-^ BMDMs in physiologic-folate media, confirming our previous ELISA data (Fig. 4C, Fig. 5C, 5D, Supp. Fig. 5C). 1C metabolism is a critical cellular source of NADPH for maintenance of cellular redox state,^10^ and, indeed, NADPH levels were decreased in *Folr2*^-/-^ cells (Fig. 5E). Consequently, there was evidence of increased ROS in *Folr2*^-/-^ BMDMs, with increased oxidized glutathione and cholesterol-5α,6α-epoxide (an oxidized form of cholesterol) and a concomitant decrease in reduced glutathione (Fig. 5C, 5F, Supp. Fig. 5C, 5D). Gene signatures associated with ROS were enriched in *Folr2*^-/-^ macrophages relative to wild type, supporting increased oxidative stress in these cells (Fig. 5G). Of note, all metabolomic differences observed between wild-type and *Folr2*^-/-^ macrophages were lost in supraphysiologic-folate media (Fig. 5D-F, Supp. Fig. 5D, 5E).

**Figure 5.**
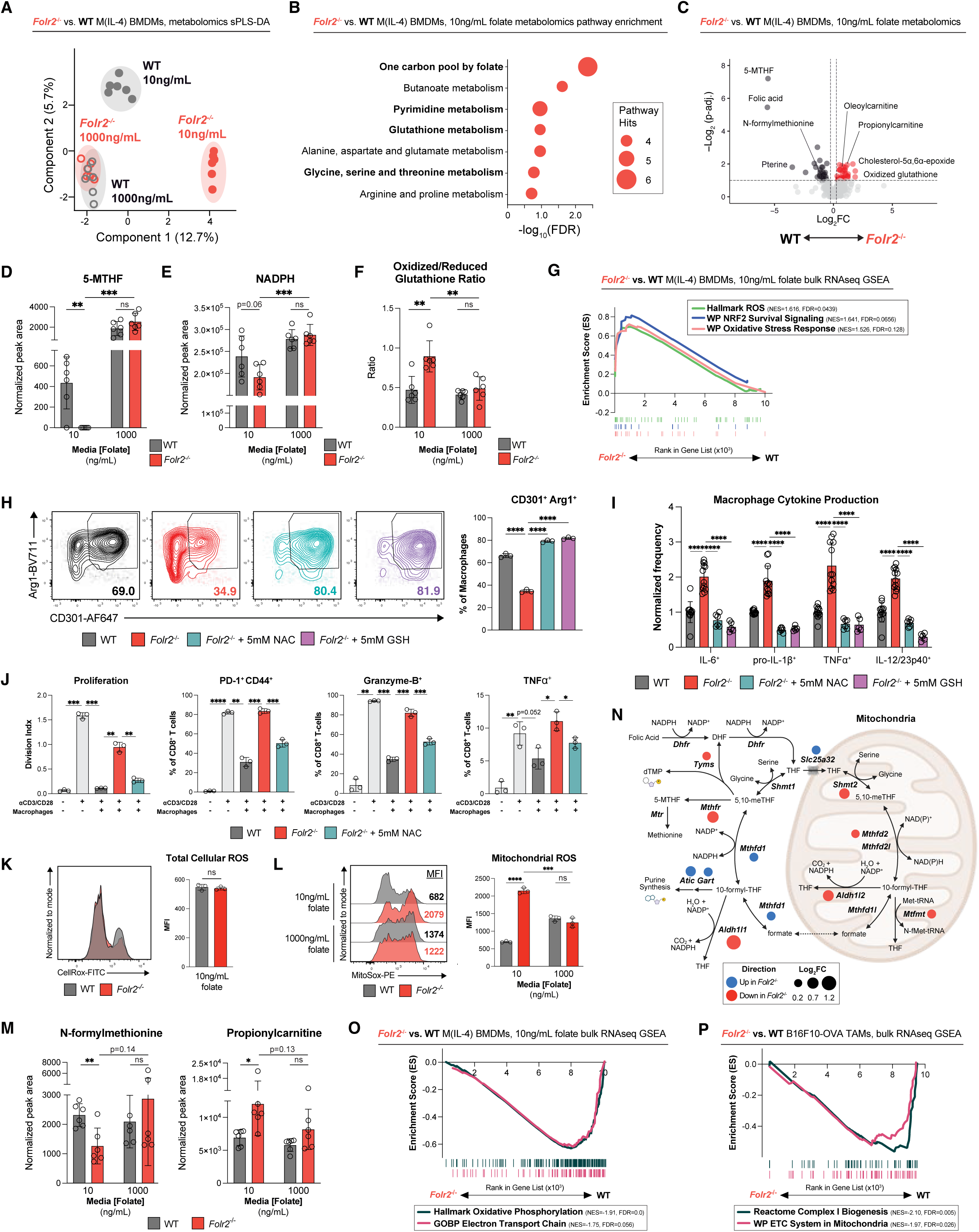
Folate deficiency induces mitochondrial oxidative stress in *Folr2*^-/-^ macrophages through dysregulation of glutathione metabolism. **(A)** sPLS-DA of untargeted metabolomics data from wild-type and *Folr2*^-/-^ M(IL-4) BMDMs grown in physiologic- or supraphysiologic-folate media. **(B)** Metabolic pathways enriched in the comparison between wild-type and *Folr2*^-/-^ BMDMs grown in physiologic-folate media. Size of dots represents the number of significant differentially detected metabolites in each pathway. **(C)** Volcano plot of differentially detected metabolites comparing wild-type and *Folr2*^-/-^ M(IL-4) BMDMs grown in physiologic-folate media. Cutoffs set at absolute Log_2_FC > 0.25, FDR < 0.1. **(D-F)** Normalized peak areas of 5-methyltetrahydrofolate (5-MTHF) **(D)** and NADPH **(E)** or reduced/oxidized glutathione ratio **(F)** detected in wild-type and *Folr2*^-/-^ M(IL-4) BMDMs grown in physiologic- or supraphysiologic-folate media. **(G)** GSEA of indicated ROS gene signatures comparing wild-type and *Folr2*^-/-^ M(IL-4) BMDMs grown in physiologic-folate media. **(H, I)** Quantification of CD301 and Arg1 expression **(H)** or intracellular cytokine production **(I)** in wild-type and *Folr2^-/-^* M(IL-4) BMDMs grown in physiologic-folate media +/- 5mM glutathione (GSH) or 5mM N-acetylcysteine (NAC). One of two independent experiments shown. **(J)** Quantification of T cell proliferation by CellTrace Violet or T cell activation markers (Granzyme-B, TNFα, PD-1^+^CD44^+^) after co-culture of CD8^+^ T cells with wild-type or *Folr2*^-/-^ M(IL-4) BMDMs at a 1:4 (BMDM:T cell) ratio. Prior to coculture, BMDMs were grown in physiologic-folate media +/- 5mM N-acetylcysteine (NAC). One of two independent experiments shown. **(K, L)** Representative flow plots and quantification of CellRox **(K)** or MitoSox **(L)** staining of wild-type and *Folr2*^-/-^ BMDMs grown in physiologic- or supraphysiologic-folate media. **(M)** Normalized peak area of N-formylmethionine and propionylcarnitine detected in wild-type and *Folr2*^-/-^ M(IL-4) BMDMs grown in physiologic- or supraphysiologic-folate media. **(N)** Schema of cytosolic and mitochondrial 1C metabolic pathway, with enzymes identified as transcriptionally upregulated (blue) or downregulated (red) in *Folr2*^-/-^relative to wild-type M(IL-4) BMDMs. Size of the colored dots represents Log_2_FC. Enzymes without dots are not differentially expressed. **(O, P)** GSEA of mitochondrial gene signatures in the ranked list of genes differentially expressed between wild-type and *Folr2^-/-^* M(IL-4) BMDMs **(O)** and TAMs flow-sorted from B16F10-OVA tumors **(P)**. * p < 0.05; ** p < 0.01; *** p < 0.001; **** p < 0.0001; ns not significant. BioRender license: Created in BioRender. Zimmerman, M. (2027) https://BioRender.com/8178nij

To test if excess ROS induces pro-inflammatory functions in *Folr2*^-/-^ BMDMs, we treated cells with glutathione (GSH) or n-acetylcysteine (NAC) to decrease ROS levels. Treatment with either antioxidant increased expression of the immunosuppressive markers CD301 and Arg1 and decreased production of IL-6, IL-12/23p40, TNFα, and pro-IL-1β by *Folr2*^-/-^ BMDMs (Fig. 5H, 5I). Furthermore, pre-treatment of *Folr2*^-/-^BMDMs with NAC prior to co-culture with activated T cells increased the T cell suppressive capacity of *Folr2*^-/-^BMDMs (Fig. 5J). While ROS have been shown to activate the inflammasome, prior work in the THP-1 leukemia cell line suggests that FRβ activates the inflammasome independent of folate.^31^ We did not observe changes in ASC puncta formation or inflammasome-mediated IL-1β cytokine secretion in *Folr2*^-/-^ BMDMs (Supp. Fig. 5F, 5G). Together, these functional studies demonstrate that excess ROS produced in *Folr2*^-/-^BMDMs are critical for the pro-inflammatory functions of *Folr2*^-/-^ BMDMs independent of the inflammasome.

Given that the location of ROS production is critical for their physiologic functions, we next interrogated ROS subcellular localization. To our surprise, we observed no difference in CellRox staining, a global cellular oxidative stress indicator, between wild-type and *Folr2*^-/-^ BMDMs in physiologic-folate media (Fig. 5K). However, there was a significant increase in MitoSox staining, a mitochondria-localized superoxide probe, in *Folr2*^-/-^ BMDMs that could be rescued by supraphysiologic-folate media (Fig. 5L), suggesting mitochondrial ROS accumulation is the source of the oxidative stress observed in *Folr2*^-/-^ BMDMs. Returning to our metabolomic data, we identified a clear pattern of mitochondrial dysfunction. N-formylmethionine was decreased in *Folr2*^-/-^ BMDMs, which suggests decreased mitochondrial translation (Fig. 5C, 5M).^49^ We also observed an increase in acylcarnitines (propionylcarnitine, tetradecanoylcarnitine, and oleoylcarnitine), which are the substrates for mitochondrial fatty acid beta-oxidation, suggesting decreased energy production from fatty acids in *Folr2*^-/-^ BMDMs (Fig. 5C, 5M). Transcriptional analysis also supported a decrease in mitochondrial function, with depletion of oxidative phosphorylation and electron transport chain gene signatures and decreased mitochondrial 1C enzymes at the individual transcript level in *Folr2*^-/-^ BMDMs (Fig. 5N, 5O). We confirmed this decrease in oxidative phosphorylation and electron transport chain transcriptional signatures in *Folr2*^-/-^ TAMs (Fig. 5P). This integrative analysis suggests that *Folr2*^-/-^ macrophages have impaired mitochondrial 1C metabolism in physiologic levels of folate resulting in oxidative stress.

### Folate uptake by FRβ is required to prevent mitochondrial dysfunction

Given our data implicating excess mitochondrial ROS in *Folr2*^-/-^ BMDMs, we wanted to test if mitochondria were dysfunctional in *Folr2*^-/-^macrophages. We observed a decrease in mitochondrial membrane potential per unit mitochondrial mass (TMRM/MTG ratio) in *Folr2*^-/-^ BMDMs, a hallmark of mitochondrial dysfunction (Fig. 6A). Using Seahorse, we discovered *Folr2*^-/-^ BMDMs had decreased basal oxygen consumption rates (OCR) and mitochondrial ATP production as well as increased basal extracellular acidification rates (ECAR), resulting in a decreased OCR/ECAR ratio (Fig. 6B, Supp. Fig. 6A). These data suggest that *Folr2*^-/-^ BMDMs have dysfunctional mitochondria with a concomitant metabolic shift towards glycolysis. Transmission electron microscopy revealed an increase in phagolysosomes, mitochondria with disorganized cristae, and mitochondria frequently located in lysosomes in *Folr2*^-/-^ BMDMs, suggesting mitochondrial degradation (Fig. 6C, Supp. Fig. 6B). We confirmed that *Folr2*^-/-^ BMDMs exhibited an increase in mitochondrial degradation by immunofluorescence staining, revealing many more colocalized mitochondria (TOM20^+^) and lysosomes (LAMP1^+^) in *Folr2*^-/-^ BMDMs (Fig. 6D, Supp. Fig. 6C).

**Figure 6.**
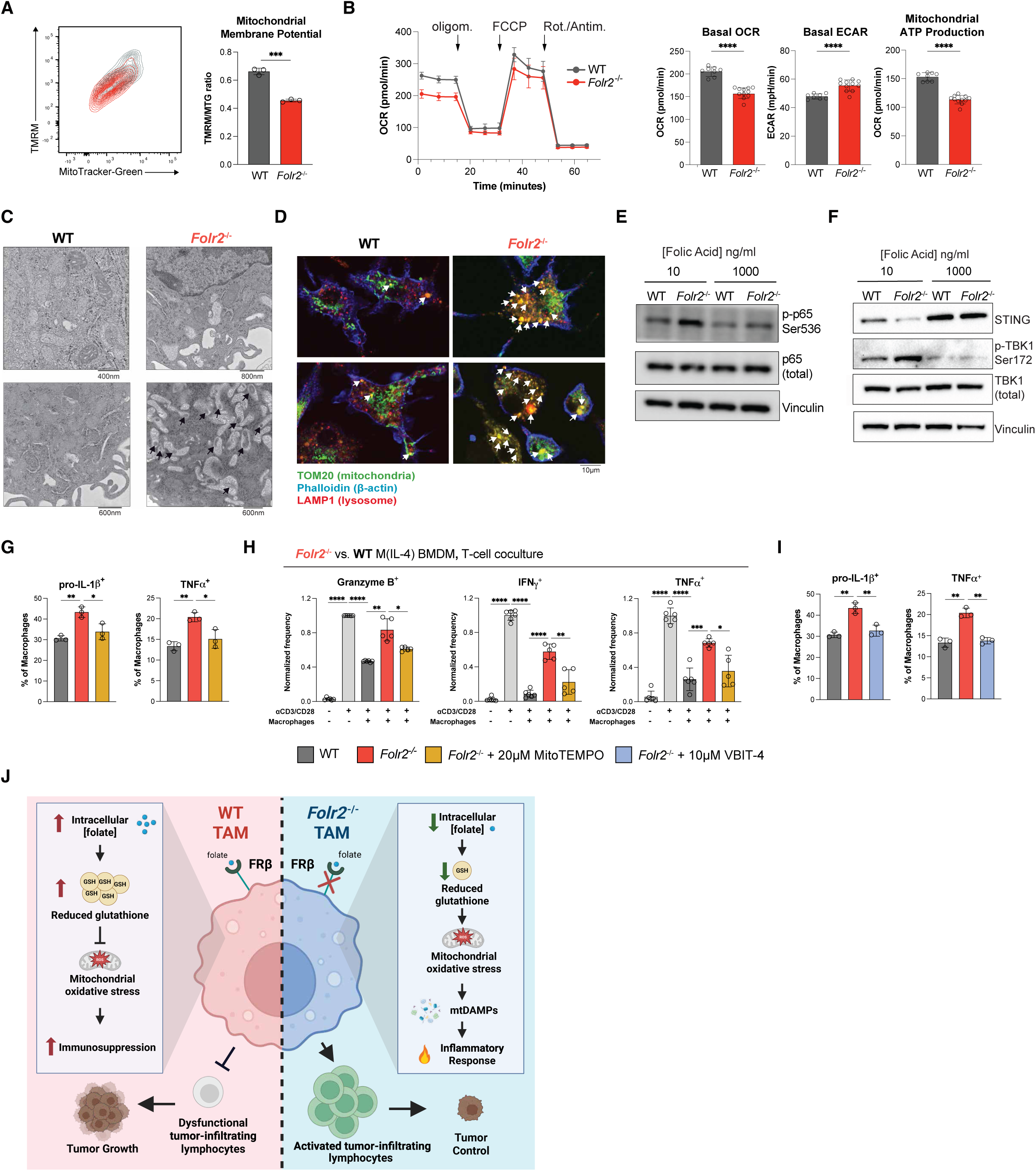
Dysfunctional mitochondria induce pro-inflammatory polarization of *Folr2*^-/-^ macrophages. **(A)** Tetramethylrhodamine methyl ester (TMRM) and Mitotracker Green staining of wild-type and *Folr2*^-/-^ M(IL-4) BMDMs grown in physiologic-folate media. Flow cytometry plot (left) and quantification of one of three representative experiments (right) shown. **(B)** Seahorse XF Mito Stress Test of wild-type and *Folr2*^-/-^ M(IL-4) BMDMs grown in physiologic-folate media. Representative curve (left) and bar plots of key measurements (right) - basal oxygen consumption rate (OCR), basal extracellular acidification rate (ECAR), and mitochondrial ATP production - shown from one of three independent experiments. **(C)** Representative transmission electron microscopy images of wild-type and *Folr2^-/-^*M(IL-4) BMDMs grown in physiologic-folate media showing mitochondria and vesicles containing mitochondrial remnants (black arrows). **(D)** Immunofluorescence images of wild-type and *Folr2*^-/-^ M(IL-4) BMDMs grown in physiologic-folate media stained with antibodies against LAMP1 (lysosomes) and TOM20 (mitochondria), as well as Phalloidin (F-actin). Mitochondria in phagolysosomes identified by overlapping staining for LAMP1 and TOM20 (white arrowheads). **(E, F)** Immunoblot of lysates from wild-type and *Folr2*^-/-^ M(IL-4) BMDMs grown in physiologic- or supraphysiologic-folate media stained against phospho/total-p65 **(E)** or STING and phospho/total-TBK1 **(F)** Vinculin used as a loading control for both **(E)** and **(F)**. **(G, I)** Quantification of intracellular cytokine production in wild-type and *Folr2*^-/-^ M(IL-4) BMDMs grown in physiologic-folate media +/- 20μM MitoTEMPO **(G)** or 10μM VBIT-4 **(I)**. One of two independent experiments shown. **(H)** Quantification of T cell intracellular cytokine production (IFNу, TNFα), and T cell activation markers (Granzyme B) after co-culture of CD8^+^ T cells with wild-type or *Folr2*^-/-^ M(IL-4) BMDMs at a 1:4 (BMDM:T cell) ratio. Prior to coculture, BMDMs were grown in physiologic-folate media +/- 20μM MitoTEMPO. Two pooled experiments shown. (J) Schema of proposed mechanism for the role of FRβ and folate metabolism in TAMs. * p < 0.05; ** p < 0.01; *** p < 0.001; **** p < 0.0001; ns not significant. Created in BioRender. Zimmerman, M. (2027) https://BioRender.com/9c5ut7w, https://BioRender.com/a508l9s

Excess ROS can directly activate pathways important for pro-inflammatory differentiation, such as NF-κB.^19,50^ However, mitochondrial ROS can also induce mitochondrial DNA release into the cytoplasm, activating cGAS-STING.^51^ Activated STING recruits the cytosolic kinase TBK1 to activate IRF3- and NF-κB-mediated signaling and induce pro-inflammatory cytokine production.^52^ Given our transcriptional data suggesting activation of canonical NF-κB signaling in *Folr2*^-/-^ TAMs and BMDMs (Supp. Fig. 2H, Fig. 4F), we assessed engagement of this pathway by immunoblot. *Folr2*^-/-^ BMDMs exhibited increased NF-κB p65 phosphorylation in physiologic-folate media that was rescued by supraphysiologic-folate media (Fig. 6E). Similarly, TBK1 phosphorylation increased and total STING levels decreased, consistent with pathway activation (Fig. 6F). We next asked if inhibition of mitochondrial ROS by the antioxidant MitoTEMPO, which selectively localizes to the mitochondria and neutralizes superoxide, could reverse the phenotype of *Folr2*^-/-^BMDMs. Indeed, treatment of *Folr2*^-/-^ BMDMs with MitoTEMPO decreased pro-inflammatory cytokine production and increased the suppressive capacity of these macrophages (Fig. 6G, 6H). Mechanistically, ROS induce mitochondrial DNA (mtDNA) release through oligomerization of VDAC1 and other channels in the outer mitochondrial membrane.^51^ To test if mtDNA leak contributes to *Folr2*^-/-^ macrophage pro-inflammatory function, we treated *Folr2*^-/-^ BMDMs with VBIT-4, a small molecular inhibitor of VDAC1 oligomerization.^51^ Treatment with VBIT-4 reduced inflammatory cytokine production by *Folr2*^-/-^ BMDMs (Fig. 6I). In summary, our data support that folate uptake by FRβ in physiologic-folate conditions, such as the TME, is critical to prevent mitochondrial dysfunction, ROS production, and mtDNA release to promote immunosuppressive TAM polarization and function (Fig. 6J).

## Discussion

Immunosuppressive TAMs promote tumor growth, inhibit immune effector cells, and confer resistance to ICI for melanoma patients, highlighting the need for new therapeutic strategies targeting TAMs. Using a novel *Folr2*^-/-^ mouse model, we demonstrated that FRβ promotes immunosuppressive TAM polarization and function in the TME. We identified that FRβ^+^ TAMs inhibit effector TIL function to enable tumor progression. FRβ was required for folate uptake by TAMs in low-folate environments, and using fresh human melanoma resections we demonstrated that the melanoma TME is depleted of folate relative to surrounding normal tissue. We found that in the absence of folate uptake by FRβ, macrophages accumulated dysfunctional mitochondria that produced excess ROS, activated STING and NF-κB signaling, and acquired pro-inflammatory functions. Finally, we showed that FRβ^+^ TAMs are associated with resistance to ICI in melanoma patients, and loss of *Folr2* synergized with anti-PD-1 immunotherapy in mice to control tumor growth. Together, these results demonstrate that FRβ is a novel metabolic checkpoint on TAMs that is critical for immunosuppressive phenotype and function.

Importantly, we could reverse the pro-inflammatory functions of *Folr2*^-/-^ BMDMs and TAMs by supplementing the cells or mice with excess folate, demonstrating that these phenotypes are dependent on cell-intrinsic folate deficiency. Moreover, our mechanistic work demonstrated that inhibition of ROS or mitochondrial DNA release also impaired the pro-inflammatory polarization of *Folr2*^-/-^ BMDMs, further supporting our folate-dependent mechanism. Although our data did not support non-folate roles of FRβ that have been postulated in the literature,^29–32^ we cannot rule out that FRβ has other functions in TAMs in addition to folate uptake. Interestingly, prior work has suggested that not all FRβ expressed by cells can bind folate, implying there may be inactive conformations of this protein.^33^ The relevance of this is unknown, but may explain non-folate mediated roles of FRβ observed in other contexts.

ROS activation of the Keap1-Nrf2 axis is thought to be critical for immunosuppressive TAM polarization.^16^ Conversely, acute production of ROS after LPS stimulation is required for optimal pro-inflammatory polarization of macrophages.^19,20^ ROS are complicated signaling molecules, with many species having very short half-lives. As a result, the location and type of ROS production, duration of ROS signaling, and local concentration of antioxidants all factor into the downstream effects of ROS on macrophage signaling and function.^14,15^ We identified that excess mitochondrial ROS were responsible for the phenotype of *Folr2*^-/-^ BMDMs. We hypothesize that impaired control of ROS is due to decreased mitochondrial 1C metabolism, which is essential for generating reducing power.^53,54^ There may also be a feed-forward loop occurring in our *Folr2*^-/-^ macrophages, where excess ROS damage the electron transport chain, resulting in increased superoxide production.^55^ We propose that excess ROS in mitochondria induces oligomerization of VDAC1 channels, allowing mitochondrial DNA escape to activate cGAS-STING, resulting in pro-inflammatory polarization of *Folr2*^-/-^ macrophages. Oxidized mtDNA release into the cytosol and subsequent STING activation have been described in LPS- and ATP-stimulated macrophages.^56^ We observed the same pathway engaged by folate restriction alone, indicating that metabolic stress is sufficient to trigger it — a route accessible to TAMs in the folate-depleted TME, where microbial inflammatory stimuli are typically absent.

Although we believe that our BMDM model system recapitulates the key findings of *Folr2*^-/-^ TAMs *in vivo* given the similar functional, phenotypic, and transcriptional changes observed, there were also key differences. First, we have not observed a decrease in *Folr2*^-/-^ TAM numbers *in vivo*, despite the increase in cell death of *Folr2*^-/-^ BMDMs *in vitro*. This may be due to compensatory influx of new monocytes into tumors to fill the niche vacated by dying *Folr2*^-/-^ TAMs. Alternatively, this may be due to the different form of folate used in our *in vitro* studies (folic acid), whereas the predominant form of folate *in vivo* is the reduced 5-methyltetrahydrofolate (5-MTHF). RFC binds folic acid with low affinity, so it is likely that the *Folr2*^-/-^ BMDMs were more folate deficient than *Folr2*^-/-^ TAMs.^57^ Finally, although our *in vivo* single-cell RNA-seq data granted us a detailed view of the immune infiltrates in *Folr2*^-/-^ mice, it did not allow us to determine which effects were directly due to loss of *Folr2* and which were secondary effects from downstream activation of other immune cells (e.g. increased IFNу production by activated T cells). For example, there were striking changes in neutrophils resulting in differentiation towards an interferon-sensing transcriptional state (Supp. Fig. 2I-O). We did not detect *Folr2* expression in tumor-associated neutrophils (Supp. Fig. 2D, 2E, 2N), nor was the protein FRβ found by flow cytometry on these cells (Fig. 1E), suggesting these changes were due to secondary effects. One of the enriched TAM cell states found in *Folr2*^-/-^ mice was efferocytotic TAMs that express high levels of the receptors *Axl* and *Mertk*, which promote immunosuppressive TAM functions. We hypothesize that these efferocytic TAMs may counteract the loss of other pro-tumor TAM states in *Folr2*^-/-^ mice and serve as a compensatory mechanism of immunosuppression.

Our data suggest that folate is a key but limited nutrient in the TME and that FRβ allows TAMs to obtain folate in this environment. Given FRβ transports folate via endocytosis instead of transporter-mediated channeling across the plasma membrane, FRβ is likely not the highest capacity transporter of folate in high folate concentrations. We speculate that only in low-folate conditions does the high affinity of FRβ for folate become important, suggesting that FRβ^+^ TAMs may have a unique vulnerability to FRβ inhibition because of the low folate concentration within the TME.

## Methods

### Mice

Wild-type C57BL/6J (strain #000664) female mice were initially purchased from The Jackson Laboratory. *Folr2*^-/-^ mice were generated on the C57BL/6J genetic background by the UNC Animal Models Core Facility by micro-injection of CRISPR/Cas9-engineered embryos into pseudo-pregnant dams. In brief, guide RNAs 5g67B (5’-GCACTCCACTCTGTATCAGC-3’) and 3g74B (5’-CGGTTCATCCACCTTCTAGG-3’) designed to introduce double-strand breaks at sites flanking the entire *Folr2* coding sequence (exons 3-6) were cloned into T7-tRNA-gRNA constructs (UNC Animal Models Core). After gRNA validation, C57BL/6J zygotes were microinjected with 400nM high-fidelity eSpCas9 protein (UNC Animal Models Core/UNC Protein Expression and Purification Core Facility), 600nM of each T7-tRNA-gRNA construct and 20ng/μl of donor vector DNA in microinjection buffer (5 mM Tris-HCl pH 7.5, 0.1 mM EDTA). Donor vector DNA contained a 984bp 5’ homology arm, the 4348bp critical region flanked on both sides by loxP sites, and a 1005bp 3’ homology arm. Injected embryos were implanted into recipient pseudo-pregnant B6D2F1 females. Resulting pups were screened by PCR and confirmed by genetic sequencing for the presence of the floxed allele (i.e. successful incorporation of donor vector DNA at double-strand break sites) as well as the deletion allele (i.e. unsuccessful incorporation of donor vector DNA at double-strand break sites). Pups were bred to generate *Folr2^-/-^* and control animals. Genotyping primers were designed outside of Exon 3 and Exon 6 such that a product was only produced under normal PCR conditions if the deletion allele was present. *Folr2* alleles were genotyped by touchdown PCR. The wild-type allele was detected with Folr2-SRF1 (5′-CTACCCTTGGTCTGGTTCAAC-3′) and 1159Folr2-3SqR1 (5′-GAAGCAACTAGATTCCTACAGGCTAC-3′), yielding a 326-bp product. The deleted allele was detected with Folr2-5ScF1 (5′-CACTGTCAGGATGGGCTGTCA-3′) and 1159Folr2-3SqR1, yielding a 285-bp product from the knockout allele (>4.6 kb from wild type). Both reactions used Taq polymerase under the following program: 95°C for 2 min; 15 cycles of 95°C for 30s, annealing for 30s beginning at 72°C and decreasing 1°C per cycle, and 72°C for 30s; 25 cycles of 95°C for 30s, 58°C for 30s, and 72°C for 30s; and a final 72°C extension for 5 min.

### Cell culture

B16F10 and MC38 melanoma cell lines were provided by Dr. Arlene Sharpe. B16F10-ova cell lines were made as described.^58^ MCA205 cells were provided by Dr. Jessica Thaxton. B16F10 and B16F10-OVA tumor cell lines were cultured in high glucose Dulbecco’s Modified Eagle Medium (DMEM) supplemented with 10% v/v fetal bovine serum (GeminiBio) and 1% v/v penicillin/streptomycin (Gibco). MC38 tumor cells were cultured in high glucose DMEM supplemented with 10% v/v fetal bovine serum (GeminiBio), 1% v/v penicillin/streptomycin (Gibco), and 1% v/v MEM-NEAA (Gibco). MCA205 tumor cells were cultured RPMI 1640 media (Gibco) supplemented with 10% v/v fetal bovine serum (GeminiBio), 1% v/v penicillin/streptomycin (Gibco), 1% v/v MEM-NEAA (Gibco), and 1mM sodium pyruvate (Gibco). YUMM1.7 cells were cultured DMEM/F-12 media (Gibco) supplemented with 10% v/v fetal bovine serum (GeminiBio), 1% v/v penicillin/streptomycin (Gibco), and 1% v/v MEM-NEAA (Gibco). The LookOut Mycoplasma PCR Detection Kit (Sigma MP0035) was used to periodically verify that all cell lines were negative for mycoplasma contamination. For BMDM cultures, C57BL/6J control or *Folr2^-/-^* mice were euthanized and whole bone marrow flushed from femurs and tibias in cold RPMI 1640 + 2% v/v fetal bovine serum and subsequently filtered through a 70μm cell strainer. RBCs were lysed with ACK lysis buffer (Gibco) and live cells counted. Cells were resuspended in folic acid-free RPMI-1640 media (Gibco) supplemented with 10% v/v dialyzed fetal bovine serum (GeminiBio 100-108), 1% v/v pen-strep (Gibco), 1% v/v MEM-NEAA (Gibco), 1mM sodium pyruvate (Gibco), 10mM HEPES (Corning), and 55μM 2-ME (Gibco) and 20ng/mL M-CSF (Peprotech) and plated in non-TC treated dishes. Folic acid (Sigma-Aldrich) was added to culture media to a final concentration of 10ng/mL (“low folate”) or 1000ng/mL (“high folate”) where indicated. Half media changes and fresh M-CSF additions were performed every 2-3 days for a total of 6-7 days in culture. Terminally differentiated macrophages were confirmed >95% CD11b^+^F4/80^+^ at this time (data not shown). For M(IL-4) BMDMs, 20ng/mL IL-4 was added for the final 24 hours of culture prior to downstream analysis. Mouse T cells were isolated from C57BL/6J spleens by CD8^+^ MACS negative selection (Miltenyi), counted, and cultured in T cell media (TCM) (RPMI-1640 (Gibco) media supplemented with 10% FBS (GeminiBio), 1% penicillin/streptomycin (Gibco), 10mM HEPES (Corning), 55μM 2-mercaptoethanol (Gibco)). TAMs and other myeloid cells were isolated from tumors using CD11b^+^ MACS beads (Miltenyi) and cultured in folate-free RPMI-1640 media (Gibco) supplemented with 10% v/v dialyzed fetal bovine serum (GeminiBio 100-108), 1% v/v pen-strep (Gibco), 1% v/v MEM-NEAA (Gibco), 1mM sodium pyruvate (Gibco), 10mM HEPES (Corning), 55μM 2-ME (Gibco), and 10ng/mL folic acid.

### *In vivo* tumor studies

Tumor cells were cultured to approximately 80% confluency, lifted from plates with 0.25% trypsin + EDTA (Gibco), and live cells counted. Cells were diluted to a concentration of 10^7^ cells/mL in 1x HBSS (Gibco) and 100uL (1×10^6^ cells) were injected subcutaneously into the flanks of male or female 8-12 week old control or *Folr2*^-/-^ mice. Mice were monitored every 2-3 days and tumor growth measured by digital calipers, with tumor volume estimated by the formula (π*(length*width^2^))/6. Mice were fed low folate diet (estimated 0.2mg/kg; Inotiv TD.95247) or high folate (8.6mg/kg folic acid; LabDiet Select Rodent 50 IF/6F Auto) chow diets for at least 4 weeks prior to tumor implantation. Mice were treated with 100μg anti-PD-1 (clone 29F.1A12, BioXCell) or isotype control antibody (clone 2A3, BioXCell) by intraperitoneal injection on the indicated days after tumor implantation. CD8^+^ T cell depletion was accomplished by treating mice with 200µg anti-CD8a (clone 2.43, BioXCell) or the appropriate isotype control antibody (clone LTF-2, BioXCell) starting the day prior to tumor implantation and treating every 2-3 days. For intratumoral BMDM transfer studies, B16F10-OVA tumor cells were cultured and implanted as above into C57BL/6J mice fed a high folate diet. Concurrently, wild-type or *Folr2^-/-^*BMDMs were cultured in 10ng/mL folic acid media for 8 days. When tumors reached approximately 100mm^3^, 1×10^6^ wild-type or *Folr2^-/-^* BMDMs were injected intra-tumorally. Mice were determined to reach a humane endpoint and euthanized if tumor volume exceeded 2000mm^3^, tumor length surpassed 20mm, or body condition score was ≤ 2 or ≥ 4.

### *Ex vivo* human and mouse tumor interstitial fluid and tumor-infiltrating leukocyte analysis

Melanoma patients undergoing metastasis resection to develop a TIL cellular product (Lifileucel) were consented to donate extra tumor tissue for research studies (IRB #90-0573, LCCC 9001 biobanking protocol and IRB #25-0865). Tumors were delivered on ice in PBS to the Miller lab immediately after collection. After blotting dry, the tumors were placed on a 20µm nylon net mesh filter over the opening of a 5mL eppendorf tube and spun (400rcf x 15min at 4°C) to collect tumor interstitial fluid, in which folate was measured by ELISA (see below). Tumors were then mechanically minced and placed in digestion media (RPMI-1640 (Gibco) + 2% FBS (GeminiBio) + 2mg/mL collagenase P (Sigma) + 50mg/mL DNase I (Sigma)) for 30 minutes at 37°C. Fully digested tumors were vortexed, filtered through a 70µm cell-strainer, and washed in cold R2 (RPMI-1640 (Gibco) + 2% v/v FBS (GeminiBio)). Immune cells were isolated using MACS (CD45^+^ TIL or CD11b^+^) bead-based positive selection (Miltenyi) to purify immune cells for flow cytometry analysis. For murine studies, after euthanasia of the mice tumors were dissected and TIF harvested. Tumors were then minced and digested as above, followed by isolation of immune cells by MACS (CD45^+^ or CD11b^+^) bead-based positive selection.

### Isolation of macrophages from mouse organs

To isolate resident tissue macrophages from mouse organs, 8-12 week old C57BL/6J mice that had been placed on folate-deficient chow for 4.5 - 5.5 weeks were euthanized and kidneys, lungs, livers, and spleens removed. For kidneys, lungs and livers, organs were mechanically minced using dissection scissors and digested in 5mL of organ dissociation media (R2 + 1mg/mL collagenase I (Sigma) + 50ug/mL DNase I (Sigma)) for 45min at 37°C, with periodic vortexing at 15min intervals. Kidney, lung and liver suspensions were pulse vortexed, filtered through a 70µm cell strainer, and RBC lysis was performed using ACK lysis buffer (Gibco). Resulting single cell suspensions were CD45+ MACS positively selected (Miltenyi). For spleen processing, organs were mashed through a 70µm cell strainer to generate a single cell suspension and RBC lysis was performed using ACK lysis buffer (Gibco). Lysed spleens were spun down, washed, and refiltered through a 70µm cell strainer.

### *In vitro* and *ex vivo* macrophage functional studies

Wild-type or *Folr2^-/-^* BMDMs were cultured as above in low or high folate media for 5-6 days, followed by 24 hours with 20ng/mL IL-4 (Peprotech). For macrophage cytokine production measurements, 1X Protein Transport Inhibitor Cocktail (eBioscience) containing Brefeldin A and Monensin was added to the wells 15-18 hours prior to harvest to sequester cytokines intracellularly. BMDMs were then lifted from the plates, counted, and extracellular and intracellular flow cytometry performed. For T cell suppression assays, CD8^+^ T cells from C57BL/6J spleens were isolated as above. Purified CD8^+^ T cells were resuspended to a concentration of 5×10^6^ cells/mL in pre-warmed PBS and labeled with Cell Trace Violet (CTV) (ThermoFisher Scientific) at a 1:1000 dilution factor for 20 minutes at 37°C. Cell staining was neutralized with 5X volume of complete TCM and 5×10^4^ stained CD8^+^ T cells were plated on non-TC treated U-bottom 96-well plates that had been pre-coated for 16-24 hours at 4°C with 1mg/mL agonistic anti-CD3 (eBioscience) and anti-CD28 (eBioscience) antibodies diluted in PBS. Different numbers of BMDMs were added into the wells for the indicated BMDM:T cell ratios. For the final 6 hours prior to T cell coculture harvest, 1X Protein Transport Inhibitor Cocktail was added to the wells. After a total of 72 hours, all cells were collected and analyzed by flow cytometry to assess CTV dilution, T cell surface phenotype, and intracellular cytokine production. Where indicated, BMDMs were pre-treated for 4-6 days with 10µM VBIT-4 (Selleck Chemicals), 5mM N-acetyl-L-cysteine (Sigma-Aldrich), 5mM L-glutathione (Sigma-Aldrich), or 20µM MitoTEMPO (Sigma-Aldrich) prior to co-culture with T cells or cytokine measurements. Finally, myeloid cells were isolated from B16F10 tumors as above and cultured for 18 hours in folic acid-free RPMI-1640 media (Gibco) supplemented with 10% v/v dialyzed fetal bovine serum (GeminiBio 100-108), 1% v/v pen-strep (Gibco), 1% v/v MEM-NEAA (Gibco), 1mM sodium pyruvate (Gibco), 10mM HEPES (Corning), 55μM 2-ME (Gibco), and 10ng/mL folic acid (Sigma-Aldrich). For the final 6 hours of culture 1X Protein Transport Inhibitor Cocktail was added to the media, then cells lifted from the plates and analyzed by flow cytometry.

### *In vitro* BMDM survival and proliferation studies

To measure BMDM survival *in vitro*, we cultured wild-type and *Folr2^-/-^* BMDMs in media containing different concentrations of folic acid for 6-8 days. Cells were lifted, stained with flow antibodies, washed, and then stained with Annexin-V-FITC (BioLegend 640906) diluted 1:200 in Annexin-V Binding Buffer (BioLegend 422201) for 15min at 4C. Prior to flow cytometry, cells were incubated with 7-AAD per the manufacturer’s instructions (BioLegend 420404). To measure proliferation *in vitro*, we added 1mM BrdU to wild-type or *Folr2^-/-^*BMDMs cultures 9 hours prior to harvest. Cells were lifted on day 2, 4, or 6 after bone marrow collection. Cells were fixed and permeabilized using the BD Pharmingen FITC BrdU Flow Kit (BD Pharmingen 559619), then stained with anti-BrdU-FITC antibodies for flow cytometry analysis.

### Flow Cytometry

BMDMs and immune cells isolated from mouse tumors or organs for flow cytometry were blocked in MACS buffer (PBS + 2% v/v fetal bovine serum + 0.4% v/v EDTA) + TruStain FcX anti-mouse CD16/32 (BioLegend) and 2% normal mouse serum (Jackson ImmunoResearch) for at least 20 minutes at 4°C. Human immune cells were blocked in MACS buffer + TruStain FcX anti-human CD16/32 (BioLegend). Samples were then stained for surface proteins in MACS buffer for 30 minutes at 4°C (Supp. Table 5). Live/Dead fixable viability dyes (ThermoFisher Scientific) were included for dead cell exclusion. Staining master mixes were diluted in MACS buffer or BD Horizon Brilliant Stain Buffer (BD) for panels that included multiple brilliant-violet conjugated antibodies. Intracellular staining was then performed as needed by fixing and permeabilizing the cells using the eBioscience IC Fixation & Permeabilization Buffer Set (ThermoFisher Scientific) according to the manufacturer instructions. Permeabilized cells were again blocked with MACS buffer + blocking reagents listed above and then stained with antibodies diluted in 1X eBioscience Permeabilization Buffer (ThermoFisher Scientific). For analysis of mitochondrial function and redox dyes, cells were stained with 100nM tetramethylrhodamine (TMRM) (ThermoFisher Scientific), 100nM MitoTracker Green (ThermoFisher Scientific), 3µM MitoSox Red (ThermoFisher Scientific), or 5µM CellRox (ThermoFisher Scientific), each diluted in PBS, for 30 minutes at 37°C followed by subsequent washing and blocking/cell surface antibody staining. Flow data was collected on LSRFortessa (BD Biosciences), FACSymphony A3 (BD Biosciences), or Aurora (Cytek Biosciences) flow cytometers after addition of CountBright Absolute Counting Beads (ThermoFisher Scientific) to quantify cell numbers. Data was analyzed on FlowJo (v10.10.0, TreeStar), with compensation performed manually using single-color control samples. Gating was assigned using fluorescence-minus one and/or isotype-matched control antibody stained samples. For mouse samples, the following gating strategies were used: kidney, liver lung and splenic macrophages were gated on Live CD45^+^CD64^+^F4/80^+^ cells. TAMs were gated on Live CD45^+^CD11b^+^Ly6G^-^Ly6C^-^F4/80^+^. Monocytes were gated on Live CD45^+^CD11b^+^Ly6G^-^Ly6C^+^F4/80^-^ cells. Neutrophils were gated on Live CD45^+^CD11b^+^Ly6G^+^Ly6C^+^ cells. T cells were gated on Live CD45^+^CD11b^-^TCR-β^+^ cells. NK cells were gated on Live CD45^+^CD11b^-^TCR-β^-^NK1.1^+^ cells. For human samples, TAMs were gated on Live CD45^+^CD64^+^CD68^+^.

### ASC puncta immunofluorescence and quantification

Wild-type and *Folr2*^-/-^ M(IL-4) BMDMs were plated on sterile glass coverslips (72230-01, Electron Microscopy Sciences) for 24 hours in 24 well plates. Cells were maintained in low or high folate media until washing and fixation in fresh 4% PFA-PBS (15710, Electron Microscopy Sciences) for 10 minutes. Cells were permeabilized with 0.1% Saponin (S4521-25G, Sigma) maintained in all subsequent buffers. Cells were blocked with 5% normal goat serum (005-000-121, Jackson ImmunoResearch) and autofluorescence quenched with 50mM NH_4_Cl. Cells were stained with anti-ASC (AG-25B-0006-C100, Adipogen, 1:100), AlexaFluor 488 anti-rabbit secondary (A32731, Invitrogen, 1:400), DAPI 405 (D3571, ThermoFisher, 1:1000), and AlexaFluor 647-conjugated Phalloidin (8940S, Cell Signaling Technology, 1:400). Specks were imaged by confocal microscopy (ZEISS LSM900). Briefly, ASC specks were quantified by randomly generating at least five tiles in ZEN blue software. ASC puncta and nuclei were counted and summed across all tiles for a given replicate. Finally, ratios of total ASC puncta to nuclei were calculated.

### Mitochondrial immunofluorescence

Wild-type and *Folr2_-/-_* M(IL-4) BMDMs were cultured in low folate media until washing and fixation. Cells were fixed in paraformaldehyde (4%) in PBS for 15 minutes and then permeabilized with TritonX-100 (0.1%) for 15 minutes. BMDMs were blocked in 3% BSA in PBS for 1 hour before staining with mouse anti-TOM20 (sc-17764, Santa Cruz) or anti-LAMP1 (9091, Cell Signaling) antibodies in 3% BSA in PBS for overnight at 4C temperature. Cells were washed with 3% BSA in PBS and incubated with Phalloidin AlexaFluor 405 (A30104, Invitrogen), goat anti-mouse IgG (H+L) AlexaFluor Plus 488 (A32723TR, Invitrogen) and goat anti-rabbit IgG (H+L) AlexaFluor Plus 647 (A32723TR, Invitrogen) in 3% BSA in PBS for 20 minutes at room temperature. Cells were imaged using the Zeiss LSM-880 confocal microscope with ZEN acquisition software (Zeiss). Image analysis was performed in Fiji ImageJ.

### Enzyme-linked immunosorbent assays

For cytokine secretion, media was harvested from wild-type or *Folr2^-/-^*M(IL-4) BMDM cultures, centrifuged at 400 rcf for 5 minutes to remove cellular contaminants, and stored at -80°C for subsequent thawing and analysis. Secreted IL-1β was quantified from generated supernatants using the Mouse IL-1 beta/IL1F2 DuoSet ELISA kit (R&D Systems) following manufacturer instructions. For intracellular folate concentration, adherent wild-type or *Folr2^-/-^*BMDMs were harvested, washed, and lysed on a nutator for 30 minutes at 4°C in Cell Lysis Buffer II (ThermoFisher Scientific) supplemented with 50mM ascorbic acid (Sigma-Aldrich) and Pierce Protease Inhibitor (ThermoFisher Scientific), with pulse vortexing at 10 minutes intervals. After cell lysis, a small aliquot of each sample was extracted for protein quantification using BradFord Protein Assay (BioRad). The remainder of each lysate was heated at 100°C for 5 minutes and subsequently centrifuged at 14000 rcf for 10 minutes at 4°C. The resulting supernatant containing extracted folate species was transferred to a chilled eppendorf tube and used to analyze intracellular folate levels using the Folic Acid ELISA Kit (CellBio Labs). To control for cell differences between samples, quantified intracellular folate levels were normalized by total protein concentrations on a per sample basis.

### Transmission electron microscopy

Macrophage samples were collected and fixed in 2.5% glutaraldehyde/2% paraformaldehyde in 0.15 M sodium phosphate buffer, pH 7.4 for 1 hour at room temperature. Cells were then pelleted and washed with 0.15 M sodium phosphate buffer pH 7.4 three times for 10 minutes each then incubated in 1% osmium tetroxide in 0.15 M sodium phosphate buffer for 1 hour at room temperature. Cell pellets were washed three times with deionized water then dehydrated through an increasing ethanol series (30%, 50%, 75%, 90%, 100%, 100%, 100%, 10 minutes each) followed by two rounds of propylene oxide for 15 minutes each. Pellets were infiltrated with a 1:1 mixture of propylene oxide:Polybed 812 epoxy resin for several hours (Polysciences, Inc., Warrington, PA) followed with incubation in 100% Polybed 812 epoxy resin overnight. Resin was exchanged once more, then allowed to cure at 60°C. Ultrathin sections (80 nm) were cut using a diamond knife on a Leica UCT7 ultramicrotome and applied to 200 mesh copper grids. Grids were stained with 4% aqueous uranyl acetate for 12 minutes followed by Reynold’s lead citrate for 8 minutes (Reynolds, 1963). Samples were viewed using a JEOL 1400 Flash transmission electron microscope operating at 80 kV (JEOL USA, Inc.) and images were acquired with an AMT NanoSprint61 ActiveVu: High Resolution CMOS TEM camera (Advanced Microscopy Techniques).

### Seahorse XF Mito Stress Test

Wild-type and *Folr2^-/-^* BMDMs were cultured in low folate media for 6 days, followed by 24 hours with IL-4, as above. Equal numbers of wild-type and *Folr2^-/-^* BMDMs (1×10^5^ cells/well) were plated onto Agilent plates coated with Cell-Tak (Corning) for at least 30 minutes prior to analysis. The Agilent Seahorse XF Cell Mito Stress Test was performed as per manufacturer’s instructions, with 12 samples per condition. The following final concentrations of mitochondrial toxins were used: 1.5μM oligomycin, 1.5μM FCCP, 2μM Rotenone + 2μM Antimycin A. Individual wells were excluded from analysis if they exhibited an outlying OCR curve, suggesting port injection failure.

### Immunoblotting

Cell lysates were prepared with RIPA Lysis and Extraction Buffer (#89900, Thermo Fisher) containing protease inhibitor cocktail (#A32953, Thermo Fisher) and phosphatase inhibitors cocktail (#4906845001, Roche). Cell lysates were kept on a rotator at 4°C for 30 minutes and centrifuged at 10,000-14,000 g for 10-12 minutes at 4°C. Protein concentrations were measured using a BCA assay kit (#23227, Thermo Fisher) following the manufacturer’s protocol. Equal amounts of total protein were separated on 4-12% or 15% Bolt Bis-Tris Plus gels (Invitrogen) and transferred at 250mA for 90 minutes at room temperature (RT) or overnight (25V for 15 hours at 4°C) onto 0.45 µm PVDF membranes (Millipore). Membranes were blocked with 5% (w/v) non-fat milk in Tris-buffered saline containing 0.05% Tween 20 (TBST) for 1 hour at RT, then incubated overnight at 4°C with primary antibodies used at 1:1000 concentration unless otherwise indicated: anti-STING (Cell Signaling, #13647), anti-phospho-STING Ser365 (Cell Signaling, #72971), anti-TBK1 (Cell Signaling, #3504), anti-phospho-TBK1 Ser172 (Cell Signaling, #5483), anti-phospho-NFkB p65 Ser536 (Cell Signaling, #3033), anti-NFkB p65 (Cell Signaling, #8242), and anti-Vinculin (1:10,000; Sigma-Aldrich, #V9131). After three 5-minute washes in TBST, membranes were incubated for 1 hour with the corresponding HRP-conjugated secondary antibodies (1:2000 anti-rabbit, Cell Signaling, #7074P2; 1:5000 anti-Mouse, GE Healthcare, Amersham Life Sciences #NA931; 1:5000 anti-Rabbit, GE Healthcare, Amersham Life Sciences #NA934). Following three 5-minute washes, protein bands were visualized and quantified by chemiluminescence using the ChemiDoc MP Imaging System (#12003154, Biorad).

### Mouse TAM and BMDM bulk RNA-seq

Wild-type and *Folr2^-/-^* M(IL-4) BMDMs were lifted and washed for RNA isolation, with separate tissue culture plates of cells used for 3 biologic replicates. For mouse TAMs, B16F10-OVA tumors were implanted into wild-type or *Folr2^-/-^* mice fed low or high folate diet. To ensure adequate cell number per sample, CD11b^+^ MACS-selected cells were randomly pooled from 3-7 individual tumors within folate diet and mouse genotype to generate a total of 3 independent biologic replicates per group. After 16 days of growth, tumors were extracted and myeloid cells isolated by CD11b^+^ MACS positive selection beads (Miltenyi), followed by flow sorting for TAMs gated as Live CD45^+^CD11b^+^Ly6G^-^Ly6C^-^F4/80^+^ cells. Total RNA was isolated from cell pellets using a Zymo RNA Extraction Kit following the manufacturer’s protocol. Total RNA quality was analyzed using an RNA ScreenTape (Agilent 5067-5576) and TapeStation 4200 (Agilent G2991AA) and the quantity was measured using an RNA Broad Range kit (Invitrogen Q10211) and Qubit Flex Fluorometer (Thermo Scientific Q33327). mRNA sequencing libraries were prepared from up to 1000ng of total RNA using a TruSeq Stranded mRNA Library Prep Kit (Illumina 20020595) following the manufacturer’s protocol (Illumina 1000000040498). mRNA library quality and quantity were measured using a D1000 DNA Screentape (Agilent 5067-5582) and a Qubit 1X dsDNA High Sensitivity Assay Kit (Thermo Fisher Q33231), combined at equal molar ratios, and denatured following the manufacturer’s protocol (Illumina 200047032). The pool was sequenced on an Illumina NextSeq2000 P3 flow cell with a 2×50bp paired-end configuration to a designed depth of 35 million clusters per library on average and cDNA generated using Illumina barcodes.

Bulk RNA-seq data were processed using Nextflow v22.04.4 (build 5706). Sequencing reads were aligned to the mouse reference genome using STAR, with the GRCm38.p3 genome assembly and the corresponding GENCODE M9/Ensembl release 84 gene annotation. Transcript-level abundance was quantified using Salmon against the corresponding reference transcriptome. The resulting expression estimates were used for downstream gene expression and differential expression analyses. For differential expression analysis, genes were retained if they were expressed at a minimum of 25 counts in all mice within at least one experimental group. Differential gene expression analysis was performed using the DESeq2 package. The resulting lists of differentially expressed genes were subsequently analyzed for gene set enrichment using Gene Set Enrichment Analysis (GSEA), including gene sets from the Hallmark, C2, and C5 collections.

### Mouse single-cell RNA-seq

B16F10 tumors were implanted into wild-type or *Folr2^-/-^*mice fed a low folate diet. On day 17 of tumor growth, tumors were extracted and all tumor-infiltrating immune cells isolated by CD45^+^ TIL MACS positive selection beads (Miltenyi) as above, followed by flow sorting for total live CD45^+^ cells. 5 mice of each genotype were treated as independent replicates and labeled with individual TotalSeq-B Anti-Mouse Hashtag Antibodies (Biolegend) then sorted separately prior to pooling of an estimated 8000 cells from each of the 10 samples (5 wild-type, 5 *Folr2^-/-^*). Pooled samples were loaded across 2 GEM-X 3’ chip wells using the Chromium Controller (10x Genomics). Feature Barcoding and Gene Expression libraries were generated per Chromium GEM-X Single Cell 3’ v4 protocol and sequenced at a 1:9 ratio (respectively) on one lane of an Illumina NovaSeq X 25B flow cell (r1: 28bp, i7: 10bp, i5: 10bp, R2: 90bp).

Data were mapped and demultiplexed using the Cell Ranger Multi Unified Runner (10x Genomics) against the GRCm39-2024-A mouse reference genome. Cells identified as singlets were subsequently used for downstream analysis. Data preprocessing, normalization, integration, and clustering were performed using the Seurat package (v. 5.4.0) in R (v. 4.4.1). We excluded cells that did not meet any of the following criteria: fewer than 1,000 counts, fewer than 300 genes, a count-to-gene ratio above 7.5, a mitochondrial count percentage above 5%, or a ribosomal count percentage above 30%. We further applied DoubletFinder and filtered out putative doublets. All samples were then merged into a single Seurat object. The resulting expression matrix contained 29,449 cells, including 15,032 KO and 14,417 WT cells.

Dimensionality reduction and clustering were performed using Seurat’s standard clustering workflow, utilizing the functions “FindVariableFeatures,” “RunPCA,” and “RunUMAP.” Where applicable, the first 30 principal components were used. Unsupervised clustering was performed using the “FindNeighbors” and “FindClusters” functions. We first identified major cell types and found that a resolution of 0.01 was sufficient to distinguish between these clusters. Cells identified as myeloid cells, lymphocytes, and neutrophils were then analyzed as individual subsets. For each subset, we tested several clustering resolutions and selected the resolution based on the correspondence of clusters with known biological markers. Differential expression analyses between clusters and to assess the effect of the knockout were performed using a Wilcoxon rank-sum test implemented through the “FindMarkers” function. The resulting lists of differentially expressed genes were further analyzed for signature enrichment using GSEA, including signatures from the Hallmark, C2, and C5 collections. Cell-level scores for several of the more closely studied signatures were calculated using the UCell package with kNN smoothing. Trajectory analysis was performed using Monocle3 on the neutrophil subset.^59^

### Human rapid warm autopsy specimens

Warm autopsies were performed as previously described.^45^ In brief, an Autopsy Authorization Form was signed by the patient or healthcare power of attorney prior to death of the patient. Rapid autopsy was performed at the University of North Carolina at Chapel Hill and tissue samples collected no longer than 8 hours from death. Samples were collected for metabolomics by flash freezing tissues and storing them at -80°C. Formalin Fixed, paraffin embedded (FFPE) samples were collected for Visium v1 spatial analysis (10x Genomics).

### Metabolomics sample preparation

Cells were prepared for metabolomics analysis according to previously described methods.^60,61^ In brief, wild-type and *Folr2^-/-^* BMDMs were cultured in low or high folate media for 5 days, followed by 24 hours with IL-4, as above. On day 6 cells were washed 2x with ice-cold PBS, then quenched and lysed in -20°C acetonitrile and ice-cold H2O. Separate plates of BMDMs were used for 6 biologic replicates. Samples were vortexed, centrifuged at 16,000g for 10 min at 4°C, and supernatants were dried by speed vac and reconstituted in a volume of 95:5 water:methanol that was proportional to protein concentration. Tissue samples were placed in tubes with ceramic beads and homogenized in a volume of isopropanol proportional to tissue weight (2 µL/mg of tissue). Supernatants were dried by speed vac and reconstituted in 100 uL of 95:5 water:methanol. Quality control study pools (QCSPs) were created by combining aliquots of each study sample. Additionally, method blanks were created by aliquoting LC-MS grade water and performing the same extraction steps as the study samples. Final extracts were analyzed by ultra high performance liquid chromatography-high resolution mass spectrometry (UHPLC-HRMS) at the Metabolomics and Exposome Laboratory at the University of North Carolina at Chapel Hill’s Nutrition Research Institute.

### UHPLC-HRMS metabolomics data acquisition

Untargeted UHPLC-HRMS metabolomic profiling was conducted on a Vanquish UHPLC system coupled to a Q Exactive HF-X Mass Spectrometer (ThermoFisher Scientific), using previously described methods.^60,61^ Samples were placed in the autosampler in a randomized sequence, with quality control samples (QCSP) and blank injections interspersed throughout the study samples, and 5 µL was injected from each sample. Metabolite separation was performed using an HSS T3 C18 column (2.1 × 100 mm, 1.7 μm, Waters Corporation) maintained at 50°C. The mobile phase consisted of water with 0.1% formic acid (v/v) as solvent A and methanol with 0.1% formic acid (v/v) as solvent B. A gradient elution was applied, beginning at 1% B and rising to 100% B over 16 minutes, followed by a 4-minute hold, all at a constant flow rate of 400 μL/min. Full-scan mass spectra were collected across a range of 70–1050 m/z in both positive and negative mode, while MS/MS spectra were obtained through data-dependent acquisition, selecting the 20 most abundant ions for fragmentation. Raw data files were processed using Progenesis QI (version 2.1, Waters Corporation, Milford, MA, USA), and peaks showing greater average intensity in blank samples than in QC samples were excluded to eliminate background noise.

### Compound identification and annotation

Metabolite annotation was performed by matching all peaks to an in-house library of reference standards and public metabolite databases, including the NIST Tandem Mass Spectral Library and the Human Metabolome Database (HMDB). A peak was considered a match when it satisfied one or more criteria: accurate mass within 5 ppm (MS), fragmentation spectra with a similarity score greater than 30% (MS/MS), or retention time falling within ±0.5 minutes of the reference value for in-house matches only. To convey the evidence basis for each match, we applied an ontology system described previously.^60,61^ In-house library matches were labeled as OL1 when matched by MS, MS/MS, and RT together; OL2a when matched by MS and RT; and OL2b when matched by MS and MS/MS. Matches from public databases were labeled PDa (MS and MS/MS), PDc (MS and isotope pattern similarity), or PDd (MS alone). Notably, this workflow cannot always resolve certain isomeric structures from one another. In positive mode, 10,446 features were detected, of which 8,419 were annotated: 133 at OL1, 291 at OL2a, 61 at OL2b, 346 at PDa, 6,693 at PDc, and 895 at PDd. In negative mode, 6,068 features were detected, of which 4,466 were annotated: 89 at OL1, 247 at OL2a, 34 at OL2b, 145 at PDa, 3,564 at PDc, and 387 at PDd. Data were normalized using a QCSP reference sample using the “normalize to all” function in Progenesis.^62^

### Metabolomics analysis

Quality control analyses were conducted using sPLS-DA to assess the separation of experimental groups, as well as pooled samples, based on all positively and negatively charged metabolite measurements following basic filtering to remove metabolites in the bottom 10% of the IQR and the bottom 5% of expression. Our imputation strategy was based on the assumption that values below the detection threshold across the entire experimental group should be imputed as 10 - a very low value, but not too small to create an artifact during further data transformation. When most measurements within a group were missing, missing values were imputed as one-fifth of the mean value calculated from the non-missing samples. The remaining missing values were imputed using the Bayesian Principal Component Analysis (BPCA) approach. The data were subsequently normalized on a per-sample basis using median normalization, followed by log10 transformation and Pareto scaling. To assess differences between experimental groups, we first filtered the data to identify the most robust metabolite measurements, prioritizing those that matched one of the following ontology levels in the specified order: OL1, OL2a, OL2b, PDA, PDC, and PDD. Data preparation and analysis were performed using R version 4.4.1 and MetaboAnalyst version 6.0. For statistical analysis, we performed Student’s t tests and estimated log fold changes (logFC). Pathway enrichment analysis was performed using the MetaboAnalyst pipeline and the hypergeometric test against KEGG pathways. The analysis included metabolites that met the nominal significance threshold (p-value < 0.05) and were annotated in the Human Metabolome Database (HMDB).

### Human single-cell analysis

Human melanoma single-cell RNA-seq datasets were downloaded (doi.org/10.48804/GSAXBN, GSE120575).^39,40^ Sample-level single cell RNA-seq profiles were normalized using SCTransform and integrated using the RPCA method implemented in the Seurat package (DOI:10.1016/j.cell.2021.04.048). The two datasets were then integrated also using the same RPCA approach. Following dimensionality reduction via PCA, the combined object was clustered with the Louvain algorithm using the top 30 principal components. Myeloid cells were identified using canonical genes and reclustered to identify myeloid subsets.

### Human spatial transcriptomic analysis

Regions of interest from FFPE-fixed metastatic melanoma tumor resections were reviewed by a melanoma pathologist based on hematoxylin and eosin (H&E) staining, with a focus on the tumor-stroma interface. Tumors with an adequate amount of melanoma (covering at least 40% of the Visium capture area), devoid of necrosis and pigment, were QCed for RNA quality (DV200 > 30%). 5µm FFPE sample sections are mounted on the Visium FFPE v1 slides, dried overnight and processed for HE staining at (UCLA Translational Pathology Core Laboratory (TPCL)) and library construction (UCLA Technology Center for Genomics & Bioinformatics (TCGB)) according to the manufacturer’s protocol. Single-cell barcoded sequencing libraries were run on an Illumina NovaSeq 6000. Raw FASTQ files along with HE scan images were processed using the Space Ranger pipeline v1.3.0 and GRCh38-2020-A reference transcriptome. The gene counts for each Visium well in each sample were used in the following downstream analyses. The transcriptomic clustering was generated by SpaceRanger using its default graph-based Louvain clustering algorithm. We calculated cell abundances within each Visium well using cell2location,^63^ based on our combined melanoma single-cell RNA-seq datasets.^39,40^

### Human survival analysis

Gene expression data and clinical metadata for Melanoma (SKCM) from The Cancer Genome Atlas (TCGA) were downloaded from the cBioPortal website. Survival analysis was performed using the Kaplan–Meier method, with survival curves estimated using the survfit function. The ratio of *FOLR2*/*CD8A* expressions was binarized into low- and high-expression groups and used as a predictor of survival. To assess the robustness of the findings to the choice of expression threshold, multiple percentile-based cutoffs were evaluated, ranging from the 20th to the 80th percentile (0.2–0.8), with samples categorized into low- and high-expression groups at each cutoff. In addition, Cox proportional hazards regression was performed using continuous, log-transformed *FOLR2* and *CD8A* expression levels as predictors of survival. Sex, age, and body weight were included as additional covariates in the multivariable Cox regression model.

### Statistical analysis

For all tumor growth experiments, at least 5 mice per group were used per experiment with at least 2-3 independent experiments performed. Experimental results were analyzed via unpaired two-sided Student’s t-tests using GraphPad Prism 10. Outlier data points in mouse experiments were identified and removed using Rout outlier tests executed with default Prism parameters. Genomics and metabolomics data were analyzed as discussed above. Statistical significance was signified with asterisks that correspond to the following p values: ns not significant (p > 0.05), * p ≤ 0.05, ** p ≤ 0.01, *** p ≤ 0.001, **** p ≤ 0.0001.

## Acknowledgements

The UNC Animal Models Core Facility (RRID:SCR_012167), UNC Microscopy Services Laboratory (RRID:SCR_017913), UNC Hooker Imaging Core Facility (RRID:SCR_029035), and UNC Flow Cytometry Core Facility (RRID:SCR_019170) are supported in part by P30 CA016086 Cancer Center Core Support Grant to the UNC Lineberger Comprehensive Cancer Center. Library preparation, high throughput sequencing, and project management was performed by the UNC Integrated Genomics Cores (RRID:SCR_022620), LCCC Translational Genomics Lab (RRID:SCR_025231), and LCCC Office of Genomics Research (RRID:SCR_028317), which are supported in part by the School of Medicine Office of Research and funding from the Lineberger Cancer Center Support Grant (P30 CA016086-49S1) and the University Cancer Research Fund. Single-cell RNA-sequencing was performed by the CGIBD Advanced Analytics Core (RRID:SCR_015613), which is funded in part by P30 DK034987. The authors would like to acknowledge the UNC Tissue Procurement Facility for procurement, processing, storage and distribution of fresh and snap frozen malignant and non-malignant tissue (including tumor tissue and adjacent or distant non-tumor tissue from the same surgery). Histopathology/Digital Pathology was performed by the Pathology Services Core at the University of North Carolina-Chapel Hill, which is supported in part by an NCI Center Core Support Grant (P30 CA016086). Colony management services were performed by the Colony Management Core within the Division of Comparative Medicine as well as the UNC Lineberger Preclinical Research Unit at the University of North Carolina at Chapel Hill which is supported in part by an NCI Center Core Support Grant (P30 CA16086). Untargeted metabolomics was provided through NC state funds to the Sumner Lab with sample preparation, data acquisition, and quality control performed by Rachel Cobel, Emily Brasseur, and Sabrina Molina. Agilent Seahorse analysis was performed with the assistance of the UNC Nutrition Obesity Research Center (NORC) Genomics and Energy Metabolism (GEM) Core supported by the National Institute Of Diabetes And Digestive And Kidney Diseases of the National Institutes of Health under Award Number P30DK056350. The content is solely the responsibility of the authors and does not necessarily represent the official views of the National Institutes of Health. The authors would also like to thank Dr. Adam Pefferle (TGL) and Gabrielle Cannon (AAC) in particular for their assistance and technical expertise.

## Declaration of generative AI and AI-assisted technologies in the manuscript preparation process

During the preparation of this work, the authors used Claude to suggest names for single-cell clusters and highlight patterns in metabolomics data. The authors reviewed and edited the output as needed and take full responsibility for the content of the published article.

## Authors’ contributions

Conception and design: MPZ, BCM

Acquisition of data: MPZ, HW, EKC, WLC, AGB, ASK, BPF, AGR, AEM, JAM, KV, KET, PCO, SM, KEH, CWA, DWO, BRR, WH, SJM

Analysis and interpretation of data: MPZ, VZ, HW, EKC, WLC, ASK, BPF, JAM, CJW, NET, RL, KLB, ASB, JPYT, BRR, SJS, SAK, WH, SJM, JET, BCM

Writing, review, and/or revision of the manuscript: MPZ, JAM, KLB, SAK, WH, JET, BCM

Administrative, technical, or material support: DWO, SJM

Study supervision: JYPT, SJS, SAK, WH, SJM, JET, BCM

---

**Supplementary Figure 1.**
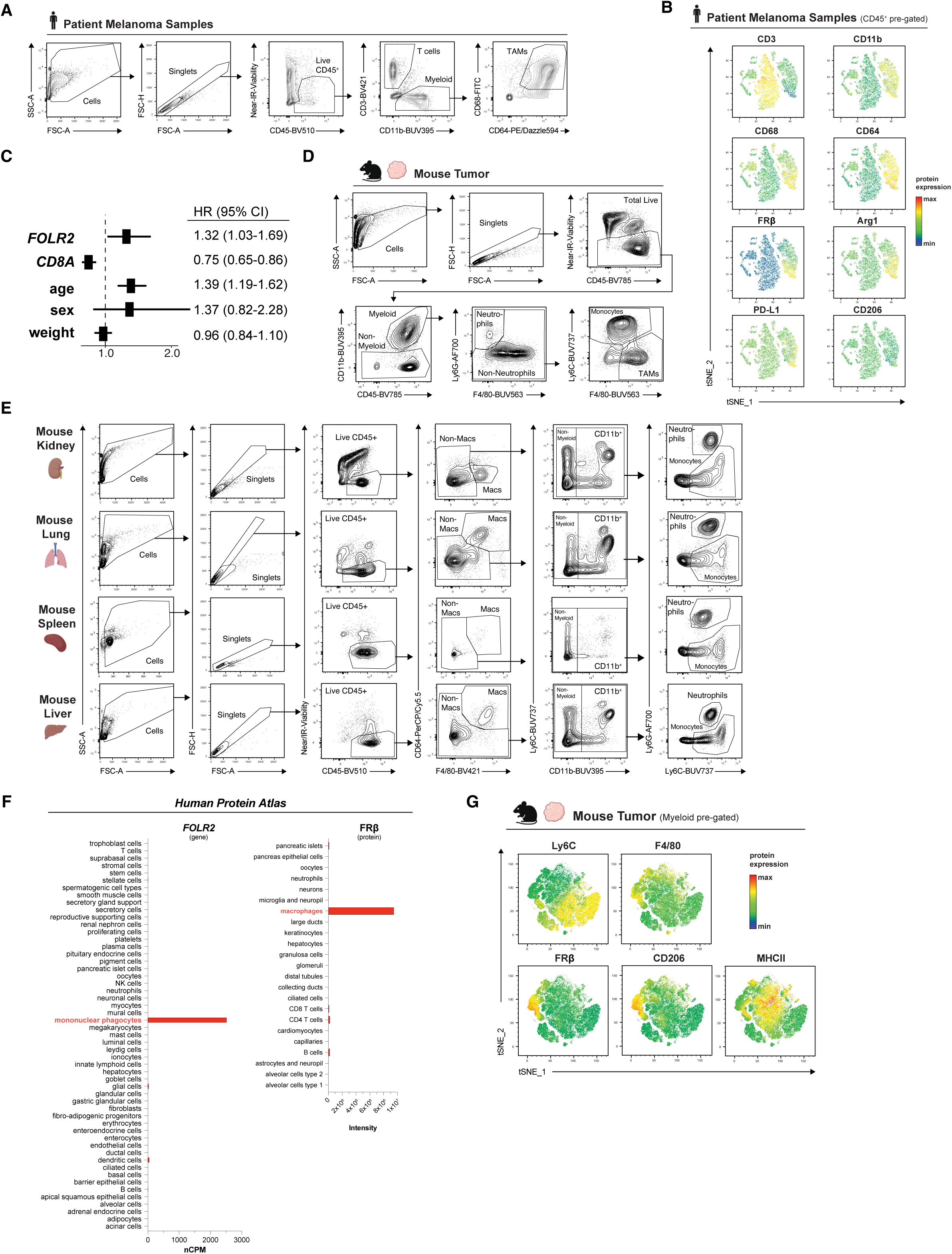
**(A)** Flow cytometry gating strategy for human patient melanoma samples. **(B)** tSNE analysis of flow cytometry data of Live CD45^+^ immune cells from one representative patient sample. **(C)** Multivariate Cox analysis of the SKCM cohort of TCGA accounting for *CD8A*, sex, age, and weight. **(D)** Myeloid flow cytometry gating strategy for mouse tumor samples. **(E)** Flow cytometry gating strategy for mouse organ samples. **(F)** *FOLR2* transcript and FRβ protein expression data from the Human Protein Atlas v25.1 (proteinatlas.org). **(G)** tSNE analysis of Live CD45^+^ immune cells from 3 pooled B16F10 tumors. Created in BioRender. Zimmerman, M. (2027) https://BioRender.com/lv1kxak

**Supplementary Figure 2.**
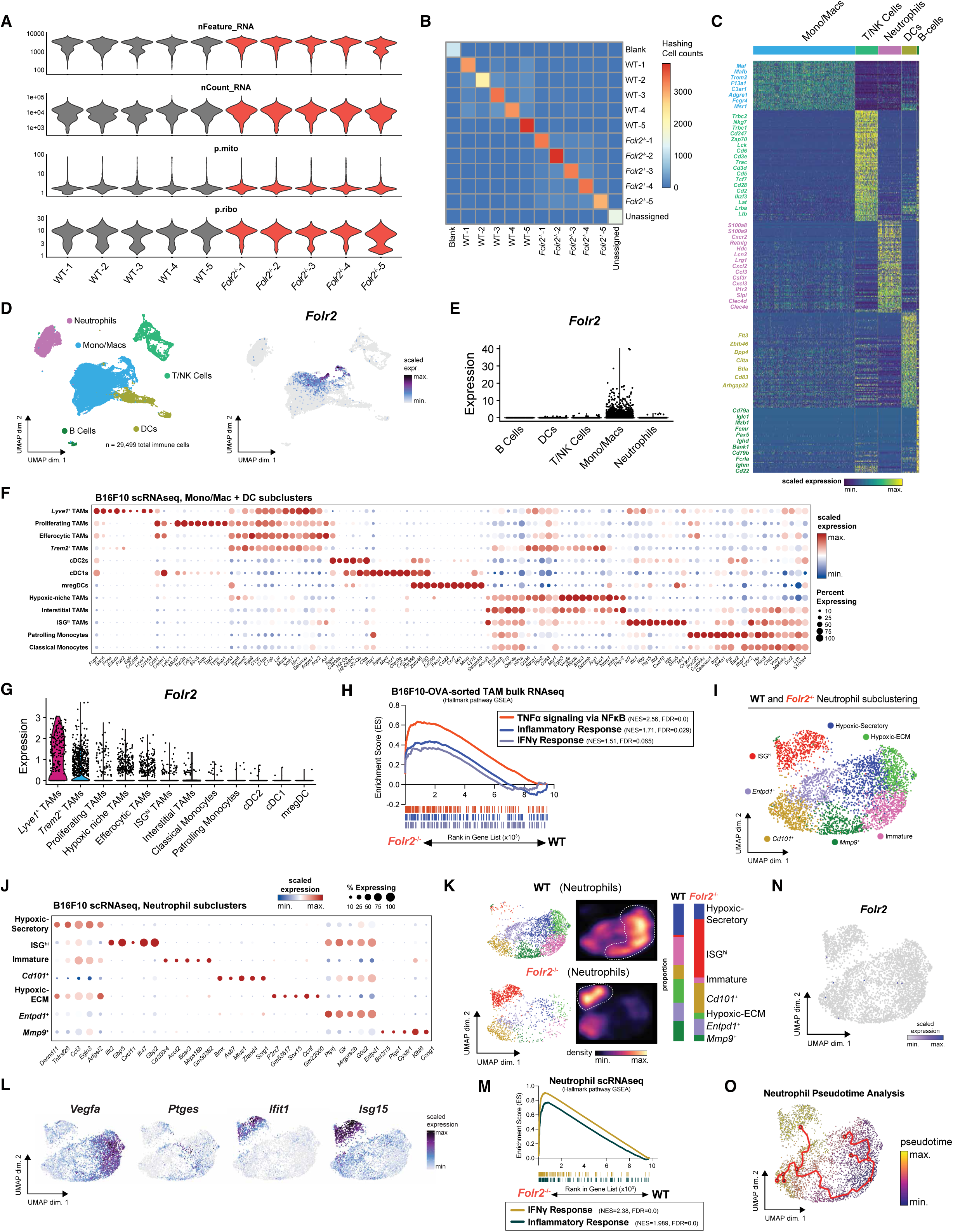
**(A)** Per sample single-cell quality metric scores. **(B)** Heatmaps demonstrating hashtag counts per sample including potential multiplets (diagonal = singlets, off diagonal = multiplets). **(C)** Heatmap of top differentially expressed genes from each major UMAP cluster. **(D, E)** Expression of *Folr2* in individual tumor-infiltrating leukocytes from B16F10 tumors. UMAP overlay **(D)** and violin plot of *Folr2* expression by major cell cluster **(E)**. **(F)** Dot plot of key representative genes for each cluster of myeloid cells from Fig. 2G. **(G)** Expression of *Folr2* in each myeloid cell cluster from Fig. 2G. **(H)** GSEA of pro-inflammatory gene signatures in the ranked list of genes differentially expressed between TAMs sorted from wild-type and *Folr2*^-/-^ mice. **(I)** scRNA-seq profiles of tumor-infiltrating neutrophils from B16F10 tumors implanted into wild-type or *Folr2*^-/-^ mice. UMAP of neutrophil clusters shown. **(J)** Dot plot of representative genes for each cluster of neutrophils from Supp. Fig. 2I. **(K)** Visualization of all cells or cell density via galaxy plot in UMAP space (left, middle) and quantification of the proportion of cells in each cluster between wild-type and *Folr2^-/-^*mice (right). **(L)** Expression of key genes in individual tumor-associated neutrophils from B16F10 tumors. **(M)** GSEA of inflammatory gene signatures in the ranked list of genes differentially expressed between neutrophils from wild-type and *Folr2*^-/-^ mice. **(N)** Expression of *Folr2* in individual tumor-associated neutrophils from B16F10 tumors. **(O)** Pseudotime trajectory analysis of neutrophil states using Monocle3.

**Supplementary Figure 3.**
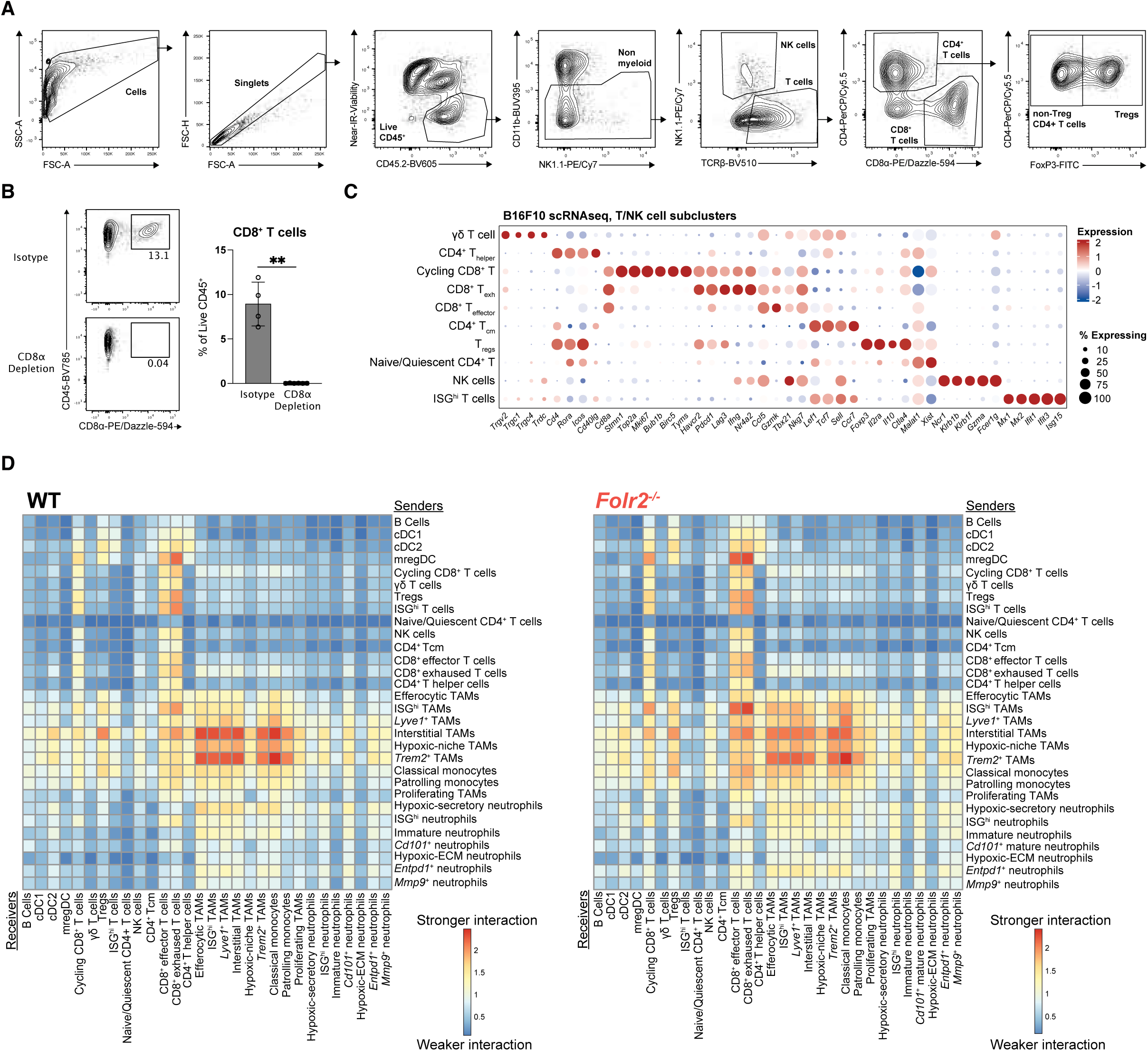
**(A)** Lymphocyte flow cytometry gating strategy for mouse tumor samples. **(B)** Representative flow plot demonstrating intratumoral CD8^+^ T cell depletion with anti-CD8α depleting compared with isotype control antibody treatment. **(C)** Dot plot of key representative genes for each cluster of lymphocytes from Fig. 3F. **(D)** Weighted interactions between clusters from CellChat analysis. Higher scores (red) indicate higher potential interactions between clusters, lower scores (blue) indicate less interaction between clusters. Separate heatmaps for wild-type and *Folr2*^-/-^ mice shown. The per cell difference between these heatmaps is plotted in Fig. 3I.

**Supplementary Figure 4.**
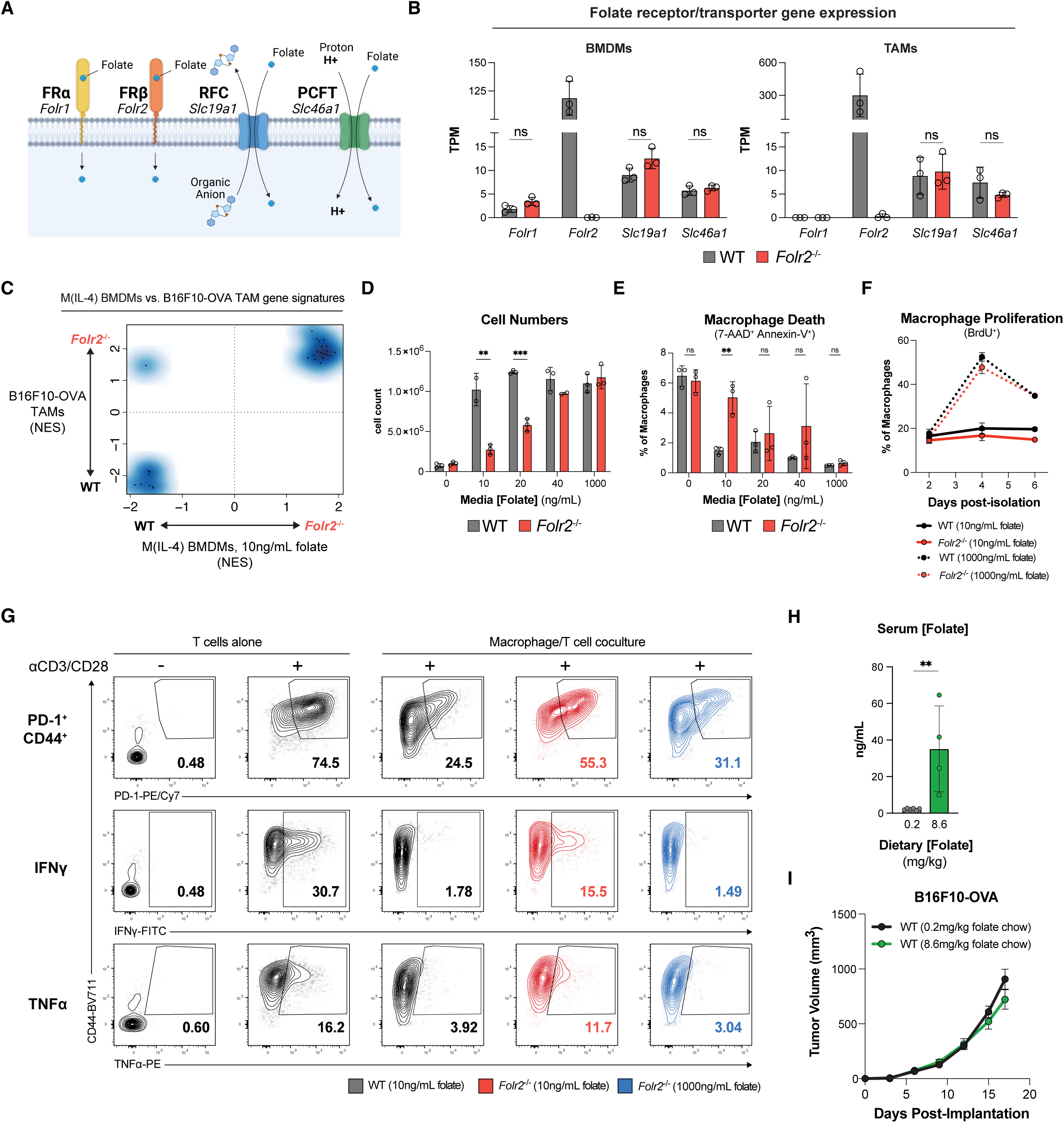
**(A)** Schema of folate receptors and transporters expressed in mice. **(B)** Folate receptor and transporter gene expression in wild-type or *Folr2*^-/-^ M(IL-4) BMDMs (left) or TAMs from B16F10-OVA tumors (right). **(C)** Plot of signatures significantly enriched (FDR < 0.1) in the comparison of wild-type vs. *Folr2*^-/-^ TAMs and wild-type vs. *Folr2*^-/-^ M(IL-4) BMDMs. Normalized enrichment score (NES) of each signature plotted on each axis. **(D)** Cell counts of wild-type or *Folr2*^-/-^ BMDMs grown for 6 days in folate-free media supplemented with the indicated concentrations of folic acid. **(E)** Flow cytometry quantification of 7-AAD^+^Annexin-V^+^ dead/dying BMDMs from experiment in **(D)**. **(F)** BrdU uptake in wild-type or *Folr2*^-/-^ BMDMs grown in 10ng/mL or 1000ng/mL folate. BrdU added to media for 9 hours prior to harvest at days 2, 4, or 6 after isolation of bone marrow from mice. **(G)** Representative flow plots of PD-1^+^CD44^+^, IFNү, or TNFα expression by CD8^+^ T cells +/- wild-type or *Folr2*^-/-^ BMDMs after *in vitro* co-culture. **(H)** Serum folate levels of mice fed a low folate or high folate diet for 4 weeks as measured by folate ELISA. **(I)** Tumor growth in wild-type mice fed low-folate or high-folate chow for four weeks prior to tumor implantation. Pooled data from 2 independent experiments shown. * p < 0.05; ** p < 0.01; *** p < 0.001; **** p < 0.0001; ns not significant. Created in BioRender. Zimmerman, M. (2027) https://BioRender.com/vbzp9l2

**Supplementary Figure 5.**
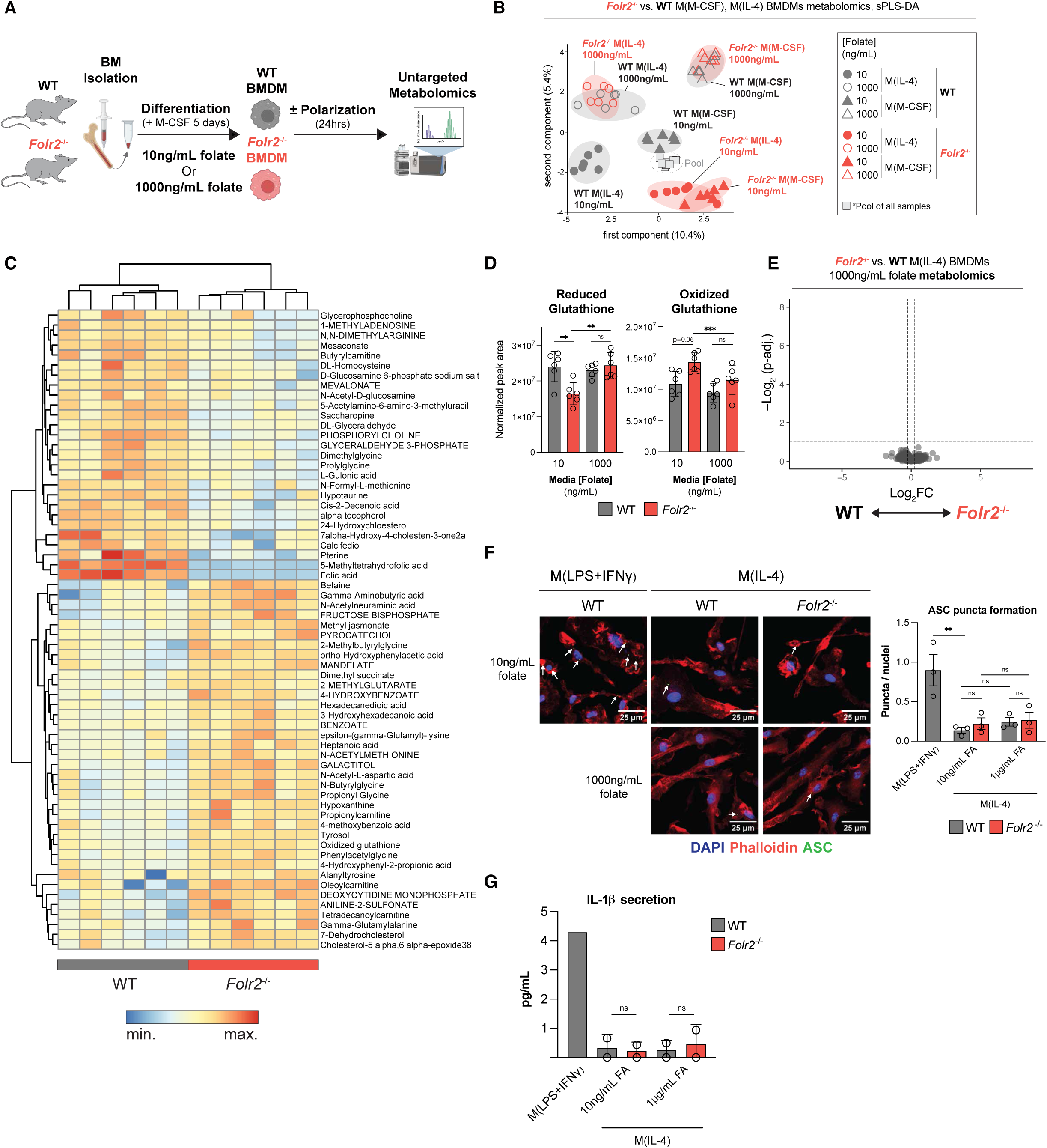
**(A)** Schema of generation of BMDMs for untargeted metabolomics. **(B)** sPLS-DA of untargeted metabolomics data from wild-type and *Folr2*^-/-^ M(IL-4) or M(M-CSF) BMDMs grow in physiologic- or supraphysiologic-folate media. **(C)** Heatmap of differentially detected metabolites (FDR<0.1) between wild-type and *Folr2*^-/-^ M(IL-4) BMDMs grown in physiologic-folate media. **(D)** Normalized peak area of reduced and oxidized glutathione detected in wild-type and *Folr2*^-/-^ M(IL-4) BMDMs grown in physiologic- or supraphysiologic-folate media. **(E)** Volcano plot of metabolites comparing wild-type and *Folr2^-/-^* M(IL-4) BMDMs grown in supraphysiologic-folate media. Cutoffs set at absolute Log_2_FC > 0.25, FDR < 0.1. **(F)** ASC puncta formation in wild-type and *Folr2*^-/-^ M(IL-4) BMDMs grown in physiologic- or supraphysiologic-folate media detected by immunofluorescence. Blue = DAPI (nuclei), Green = ASC, Red = phalloidin (F-actin). Representative images (left) and quantification (right). Wild-type BMDMs cultures in 100ng/mL LPS + 20ng/mL IFNу shown as a positive control for ASC puncta formation. White arrows identify individual puncta. **(G)** IL-1β secretion measured by ELISA from wild-type and *Folr2*^-/-^ M(IL-4) BMDMs grown in physiologic- or supraphysiologic-folate media, with M(LPS+IFNу) as a positive control for inflammasome activation. Results shown are averages from triplicate wells across 2 independent experiments. * p < 0.05; ** p < 0.01; *** p < 0.001; **** p < 0.0001; ns not significant. Created in BioRender. Zimmerman, M. (2027) https://BioRender.com/g7hgnxa

**Supplementary Figure 6.**
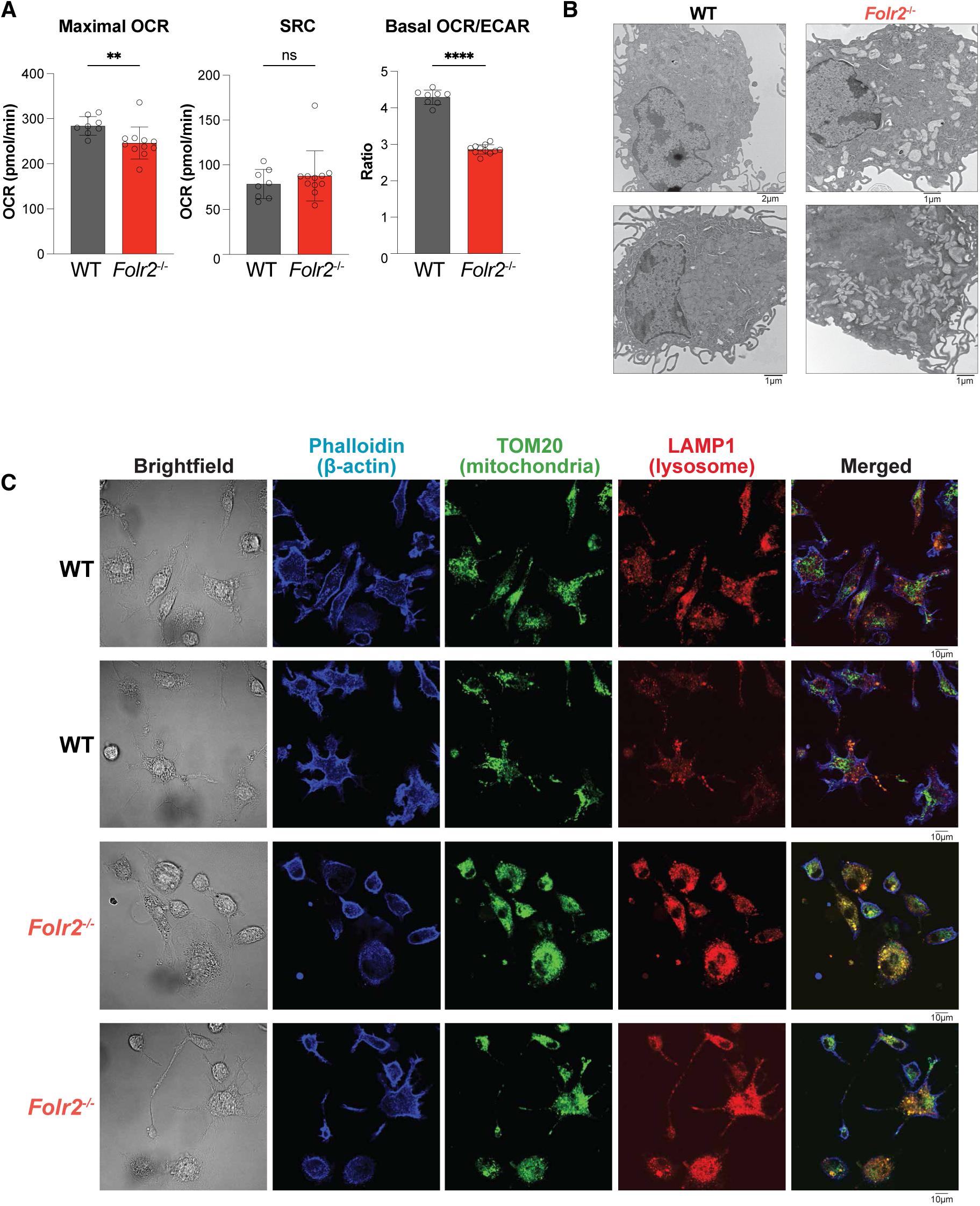
**(A)** Quantification of maximal oxygen consumption rate (OCR), spare respiratory capacity (SRC), and basal OCR/ECAR ratio from Seahorse experiment shown in Fig. 6B. **(B)** Representative transmission electron microscopy images (lower magnification) of wild-type and *Folr2*^-/-^ M(IL-4) BMDMs grown in physiologic-folate media. Same cells as shown in Fig. 6C. **(C)** Low magnification single channel and merged immunofluorescence images from cells shown in Fig. 6D.

